# Rapid emergence of latent knowledge in the sensory cortex drives learning

**DOI:** 10.1101/2024.06.10.597946

**Authors:** Céline Drieu, Ziyi Zhu, Ziyun Wang, Kylie Fuller, Aaron Wang, Sarah Elnozahy, Kishore Kuchibhotla

## Abstract

Rapid learning confers significant advantages to animals in ecological environments. Despite the need for speed, animals appear to only slowly learn to associate rewarded actions with predictive cues^1–4^. This slow learning is thought to be supported by a gradual expansion of predictive cue representation in the sensory cortex^2,5^. However, evidence is growing that animals learn more rapidly than classical performance measures suggest^6–8^, challenging the prevailing model of sensory cortical plasticity. Here, we investigated the relationship between learning and sensory cortical representations. We trained mice on an auditory go/no-go task that dissociated the rapid acquisition of task contingencies (learning) from its slower expression (performance) ^7^. Optogenetic silencing demonstrated that the auditory cortex (AC) drives both rapid learning and slower performance gains but becomes dispensable at expert. Rather than enhancement or expansion of cue representations^9^, two-photon calcium imaging of AC excitatory neurons throughout learning revealed two higher-order signals that were causal to learning and performance. First, a reward prediction (RP) signal emerged rapidly within tens of trials, was present after action-related errors only early in training, and faded at expert levels. Strikingly, silencing at the time of the RP signal impaired rapid learning, suggesting it serves an associative and teaching role. Second, a distinct cell ensemble encoded and controlled licking suppression that drove the slower performance improvements. These two ensembles were spatially clustered but uncoupled from underlying sensory representations, indicating a higher-order functional segregation within AC. Our results reveal that the sensory cortex manifests higher-order computations that separably drive rapid learning and slower performance improvements, reshaping our understanding of the fundamental role of the sensory cortex.

Despite the value of rapid learning in ecological environments, most laboratory models of rodent learning show that linking sensory cues with reinforced actions is a slow, gradual process^1–4,10^. An alternative view suggests that animals, including humans, rapidly infer relationships between cues, actions, and reinforcement (i.e. learning)^6^ even if they continue to make ongoing performance errors ^7,8,11^. Recent behavioral studies in rodents have begun to reconcile these views, arguing that latent task knowledge (i.e. discriminative contingencies) can emerge rapidly even though behavioral performance appears to improve only gradually^7^. How are these two dissociable behavioral processes—rapid acquisition of contingencies versus slower performance improvements—implemented in the brain?

An attractive brain region to consider is the sensory cortex as it is thought to subserve instrumental learning by enhancing or attenuating the representation of sensory cues that drive behavior. Plasticity of cue-related responses in the sensory cortex is thought to subserve learning as it mirrors the slow and gradual improvements in behavioral performance ^1,2,5,10^. This raises a fundamental challenge: if animals learn discriminative contingencies rapidly but cue representations in the sensory cortex change slowly^1,2,9^, the causal model linking cue-related plasticity to learning becomes problematic. One possible solution is that the sensory cortex plays a role beyond cue-related representational plasticity and directly represents high-order signals that associate reinforced actions with predictive cues. Here we focus on the auditory cortex (AC) and asked whether and how it plays a higher-order role in cue-guided learning.

We trained head-fixed, water-restricted mice to lick to a target tone (S+) for water reward and to withhold licking to a foil tone (S*−*) to avoid a timeout (auditory go/no-go task, Fig. 1a). We used simple pure tones to prevent the AC from being recruited for complex sensory processing. To confirm this, two-photon imaging of AC excitatory neurons showed that stimulus identity could accurately be decoded from AC activity from the first training day with no subsequent improvement throughout training (Supplementary Figure 1), suggesting that the AC was indeed not needed for perceptual sharpening in the task and thereby allowing us to identify possible associative functions. Performance was evaluated in each session in reinforced and non-reinforced (‘probe’) trials (Fig. 1b). Performance in probe trials revealed a rapid acquisition of task contingency knowledge which was only expressed much later in reinforced trials (Fig. 1c)^7^. Reinforcement feedback, although critical for learning, paradoxically masked the underlying task knowledge. By combining this behavioral procedure with optogenetics and longitudinal two-photon imaging, we aimed to determine how quickly animals learn stimulus-action contingencies and to define the fundamental role of the auditory cortex in sound-guided learning.

Fig. 1.
Auditory cortex silencing impairs sound-guided learning and performance during learning.
**a**, Head-fixed mice were trained on an auditory go/no-go task with ^3^ -spaced pure tones. H: hit, M: miss, FA: false alarm, CR: correct reject. **b**, Every day during training, task knowledge is probed by omitting reinforcement for 20 trials. **c**, Two distinct learning trajectories are revealed: a fast acquisition of task contingencies (measured in probe trials; green) and a slower knowledge expression (measured in reinforced trials; black). **d**, Probabilistic optogenetic silencing of the auditory cortex over learning. **e**, Testing conditions. **f**, Accuracy in reinforced light-on trials (two-way ANOVA, *p <* 10*^−^*^8^). **g**, Action rate in reinforced light-on trials (HIT, *p* = 0.07; FA, *p <* 10*^−^*^33^). See also Supplementary Figure 4. **h**, Accuracy in probe light-off trials (two-way ANOVA, *p <* 10*^−^*^4^). **i**, Tone response index in S+ trials (see Methods; two-way ANOVA, *p <* 10*^−^*^101^). Black and gray lines are individual mice and dots indicate change points (see Methods). **j**, Maximal difference between hit and FA rates in probe light-off trials over the first 6 days (t-test, *p <* 10*^−^*^3^). **k**, Hit lick latency in probe light-off trials (median *±* s.e.median; Wilcoxon test, *p* = 0.007). **l**, Accuracy in reinforced light-off trials (two-way ANOVA, *p <* 10*^−^*^8^). **m**, Action rate in reinforced light-off trials (two-way ANOVA, HIT: *p* = 0.57, FA: *p <* 10*^−^*^8^). **n**, Accuracy in reinforced light-off trials with inter-subject alignment to the day where probe accuracy *≥* 0.65 (green triangle) (two-way ANOVA, *p <* 10*^−^*^5^). Supplementary Figure 3a-c. **o**, Comparison of light-off versus light-on trials to measure auditory cortex silencing effect on on-line performance. **p**, Session density plot of accuracy in reinforced light-on against light-off. Top, control; bottom, PV-ChR2. See also Supplementary Figure 3d-g. **q**, Within subject accuracy difference in reinforced light-on and light-off trials, aligned to the day where FA rate *<* 0.3 in reinforced light-off (two-way ANOVA, *p <* 10*^−^*^15^). **r**, Within subject accuracy difference in reinforced light-on and light-off when silencing started at expert level (*n* = 4; t-test, *p* = 0.58). See also Supplementary Figure 6. mean *±* s.e.m.; \**p <* 0.05; \*\**p <* 0.01; \*\*\**p <* 0.001, n.s.: not significant.

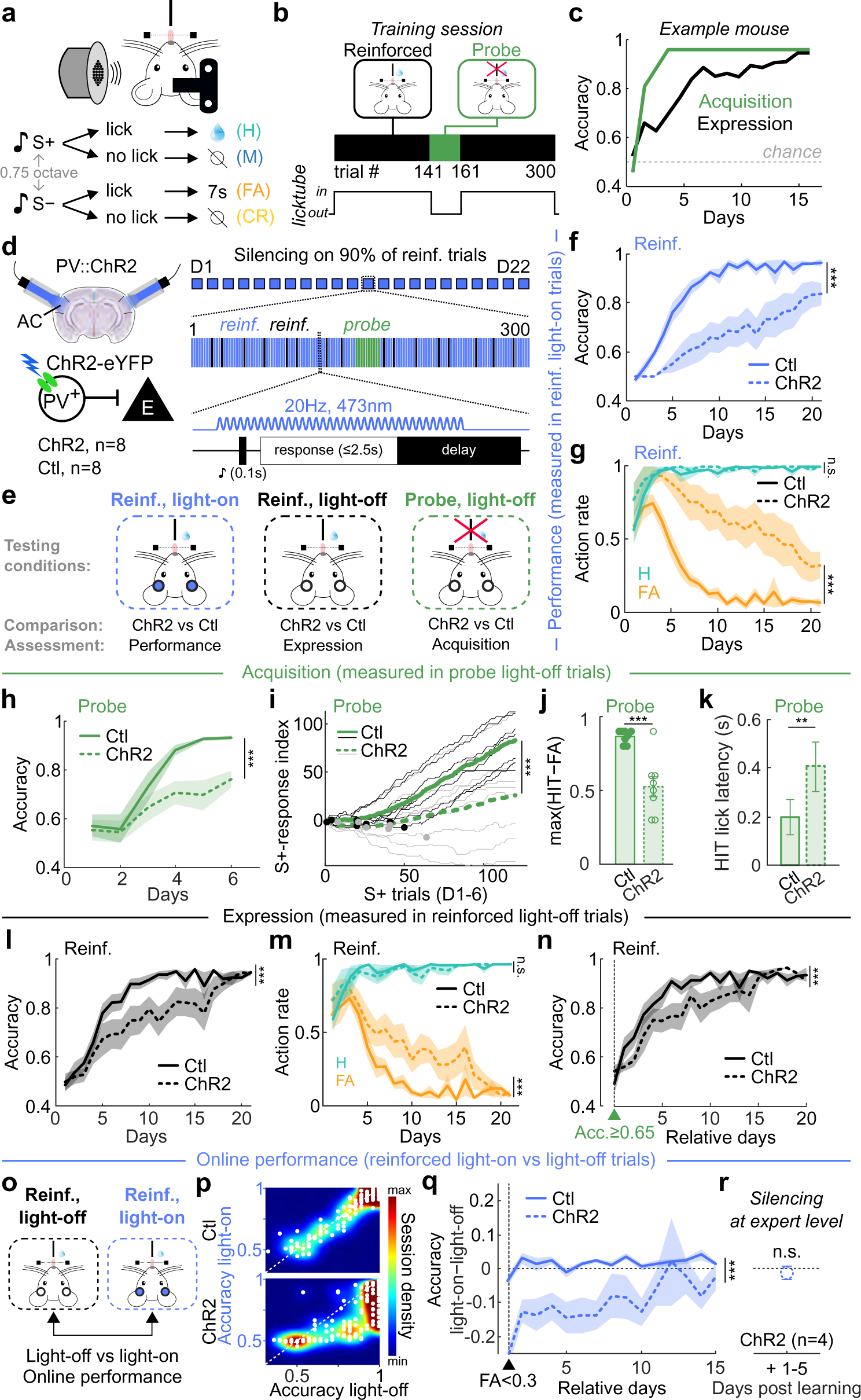

**The auditory cortex is the default pathway for sound-guided learning:** Lesion studies have suggested that the AC may not be *essential* to learn or execute cue-guided tasks with simple sensory stimuli^12–15^. However, permanent lesions cannot determine whether the AC is normally *used* for, or causally produces^16^, learning in an intact brain. To address this, we exploited a transient silencing approach to prevent the recruitment of alternative pathways^15,17–20^ while also using a probabilistic design to allow assessment of *learning* as distinct from *performance* by measuring behavior on non-silenced trials, thereby avoiding direct effects of silencing on performance.

We examined the impact of bilateral cortical silencing of the AC throughout learning (Fig. 1a). We probabilistically silenced the AC on 90% of reinforced trials throughout learning (‘light-on reinforced’, Fig. 1d), leaving 10% of reinforced (‘light-off reinforced’) and 100% of probe trials (‘light-off probe’) with intact AC activity. Silenced trials were pseudo-randomly sequenced and equally split between S+ and S*−*. Silencing was achieved by shining blue light bilaterally through cranial windows implanted above the AC of double transgenic mice (n=8) expressing channel rhodopsin (ChR2) in parvalbumin (PV) interneurons^14,21^ (Fig. 1d). We confirmed that the excitatory network was effectively silenced using this approach by combining two-photon calcium imaging of excitatory neurons and full-field optogenetic stimulation in PV-ChR2 mice (Supplementary Figure 2). Control mice (n=8) received the same light stimulation but did not express ChR2. This experimental design allowed us to assay the impact of cortical silencing on performance (control vs PV-ChR2 performance on light-on reinforced trials) versus acquisition learning (control vs PV-ChR2 performance on light-off probe trials) and expression learning (control vs PV-ChR2 performance on light-off reinforced trials) (Fig. 1e).

We first compared performance in light-on reinforced trials between PV-ChR2 and control mice (Fig. 1e) and observed a large performance impairment in PV-ChR2 mice (Fig. 1f,g). To address whether this performance reduction was accompanied by an impairment in rapid learning, we compared performance in PV-ChR2 and control animals in light-off probe trials (Fig. 1e,h-k) when the AC was not silenced and knowledge acquisition can be accurately measured^7^. Accuracy was lower during probe trials in PV-ChR2 mice (Fig. 1h), with delayed S+-response learning (Fig. 1i), lower discrimination (Fig. 1j), and longer lick latency on hit trials (Fig. 1k). Rapid acquisition of task knowledge was therefore impaired in PV-ChR2 mice.

Accuracy was also lower in reinforced light-off trials in PV-ChR2 mice (Fig. 1l,m). This remained true even after controlling for their slower task acquisition (Figs.1n, Supplementary Figure 3a-c). These impairments were also apparent in response latency and response vigor (Supplementary Figure 4). Together, these results suggest that the AC is the default pathway for sound-guided reward learning, even when not needed for perceptual sharpening.

**The auditory cortex is used during learning but becomes dispensable at expert levels:** We next sought to understand the contribution of AC activity for the expression of the learned behavior as animals transitioned to expert performance. Transient inactivation of auditory cortex in expert animals has led to conflicting results, with some reports showing degradation of sound-guided behavior^14,17,22,23^ and others not^14,24,25^. We exploited our probabilistic silencing strategy and compared performance in light-on (AC silenced) versus light-off (AC functional) reinforced trials within subjects (Fig. 1o). Performance on these two trial types was similar at early periods of training, as performance was poor overall (Fig. 1p). As training progressed, performance remained poor on light-on trials but improved on light-off trials (Fig. 1p), demonstrating that the AC is used for task performance at early and intermediate time-point during learning. Surprisingly, this deficit in performance on light-on trials gradually waned (Fig. 1p,q), suggesting that while the AC was used during learning, it became dispensable once the mice had mastered the task.

These results could be explained by three alternative explanations. First, the optogenetic manipulation *per se* may not be interfering with a task-relevant process but instead could be ‘distracting’ the animal, necessitating more time to increase performance in light-on trials. We reasoned that bilateral silencing of another cortical region that is nominally unrelated to the task would serve as an important control. We bilaterally silenced the visual cortex throughout learning in PV-ChR2 mice and found no evidence of performance impairment in light-on trials (Supplementary Figure 5), demonstrating that the performance impairment was specific to AC silencing. Second, it is possible that AC silencing altered tone perception, increasing task difficulty at the perceptual level in light-on trials. Third, the reduction of impairment during light-on trials could be driven by a reduction of the silencing effect with time due, for example, to brain damage induced by repeated silencing. To address the second and third possibilities, we trained a separate cohort of PV-ChR2 mice without daily inactivation and, instead, inactivated the AC only after they reached expert performance (see Methods). We observed no impact from AC silencing (Figs.1r, Supplementary Figure 6)^14^.

Altogether, these results show that the AC is engaged during learning but is dispensable at expert levels, potentially tutoring subcortical structures that take over once the associations are learned.

**Unsupervised discovery of learning-related dynamics by low-rank tensor decomposition:** We next sought to understand the nature and dynamics of auditory cortical activity underlying learning and performance. To do so, we performed longitudinal, two-photon calcium imaging of thousands of excitatory neurons in mice learning the auditory go/no-go task (*n* = 5). A separate group of water-restricted mice was passively exposed to two pure tones over the same duration but with no association with reinforcement (*n* = 3, see Methods; Supplementary Figure 7). This design allowed us to use the passive network as a base-case model to isolate learning-related neural dynamics.

We expressed the genetically encoded calcium indicator GCaMP6f under the CaMKII promoter, targeting AC layer 2/3 pyramidal neurons. We imaged two planes *∼*50*µm* apart (Fig. 2a), allowing us to record simultaneously hundreds of neurons per animal (n=7,137 neurons in 8 mice). All mice were passively presented with a series of pure tones (4 to 64kHz, quarter-octave spaced) to characterize auditory tuning properties within the local area of expression. We computed single-neuron tuning curves and then constructed a ‘best frequency’ map confirming the location in the AC (Fig. 2b). For each mouse, we chose two stimuli that were similarly represented in the recorded population and were 3/4 octaves apart (Fig. 2c). We used a custom head-fixation system that allowed for kinematic registration and tracked the activity of the same neurons across weeks, including pre- and post-learning tuning curve sessions (*n* = 4, 643 neurons in 8 mice, see Methods; Fig. 2d-g).

Fig. 2.
Low-rank tensor decomposition reveals learning-related network dynamics.
**a**, Multi-plane, longitudinal two-photon calcium imaging of layer 2/3 excitatory network in the auditory cortex during learning (*n* = 5 mice) or passive exposure (*n* = 3 mice; see Methods). **b**, Tonotopic organization of the field of view of one example mouse before learning (left). Cells are colored according to their best frequency and tone-evoked responses of example cells circled in black to 17 pure tones ranging from 4 to 64 kHz are displayed on the right. **c**, Tone-evoked activity (top) and proportion of responsive cells (bottom) to pure tones. S+ and S*−* (filled and unfilled triangles, respectively) are chosen for training in the task based on their equal representation in the field of view in **b**. **d**, Six example cells tracked everyday across weeks. **e**, Two planes recorded in one example mouse. Cells are colored according to the number of days tracked among the 19 recording sessions in this mouse. **f**, Distribution of number of tracked days per cells in **e**. **g**, Cumulative distribution of tracked cells according to the percentage of recording sessions. Data for mouse in **e** is the light blue line. **h**, Calcium data is arranged by neurons *×* time within trial (*−*1 to +4s relative to tone onset, vertical line) *×* trials over time *×* trial outcomes. **i**, Activity from all Learning and Passive cells are concatenated together to create a fourth-order tensor (megamouse; left). In the 3^rd^, ‘across trials’ dimension, data is aligned across mice according to learning phases: Acquisition (performance increases in probe trials), Expression (performance increases in reinforced trials), and Expert (high, stable performance in reinforced trials; see Methods and Supplementary Figure 8). **j**, Megamouse tensor decomposition identifies six neuronal dynamics (numbered; see Methods) that are characterized by a set of four factors: Neuron, Within trial, Across trial, and Outcome (see also Supplementary Figure 10). **k**, Projection of the tensor decomposition output onto principal subspace. *W*_N_*_r_*, *W*_W_ *_r_* and *W*_A_*_r_* indicate neuronal, within trial and across trial weights for a component *r*, respectively. **l**, t-distributed stochastic neighbor embedding (t-SNE) projections of neuronal weights. Each dot represents a cell, colored according to the neuronal dynamic it contributed in the most. Bars (right) display the proportion of learning and passive cells among the highest contributors for each dynamic. Dynamics 1 and 2 are driven by the passive network (burgundy), while Dynamics 3 to 6 are driven by the learning network (blue). **m**, In the passive network, the highest contributing cells in Dynamic 1 define cell ensemble 1, and highest contributing cells in Dynamic 2 define cell ensemble 2. Similarly, in the learning network, cell ensembles 3 to 6 are constituted of the highest contributing cells to Dynamics 3 to 6, respectively. **n**, Absolute weights of cell ensembles across the six identified dynamics. Neurons can participate in more than one dynamic.

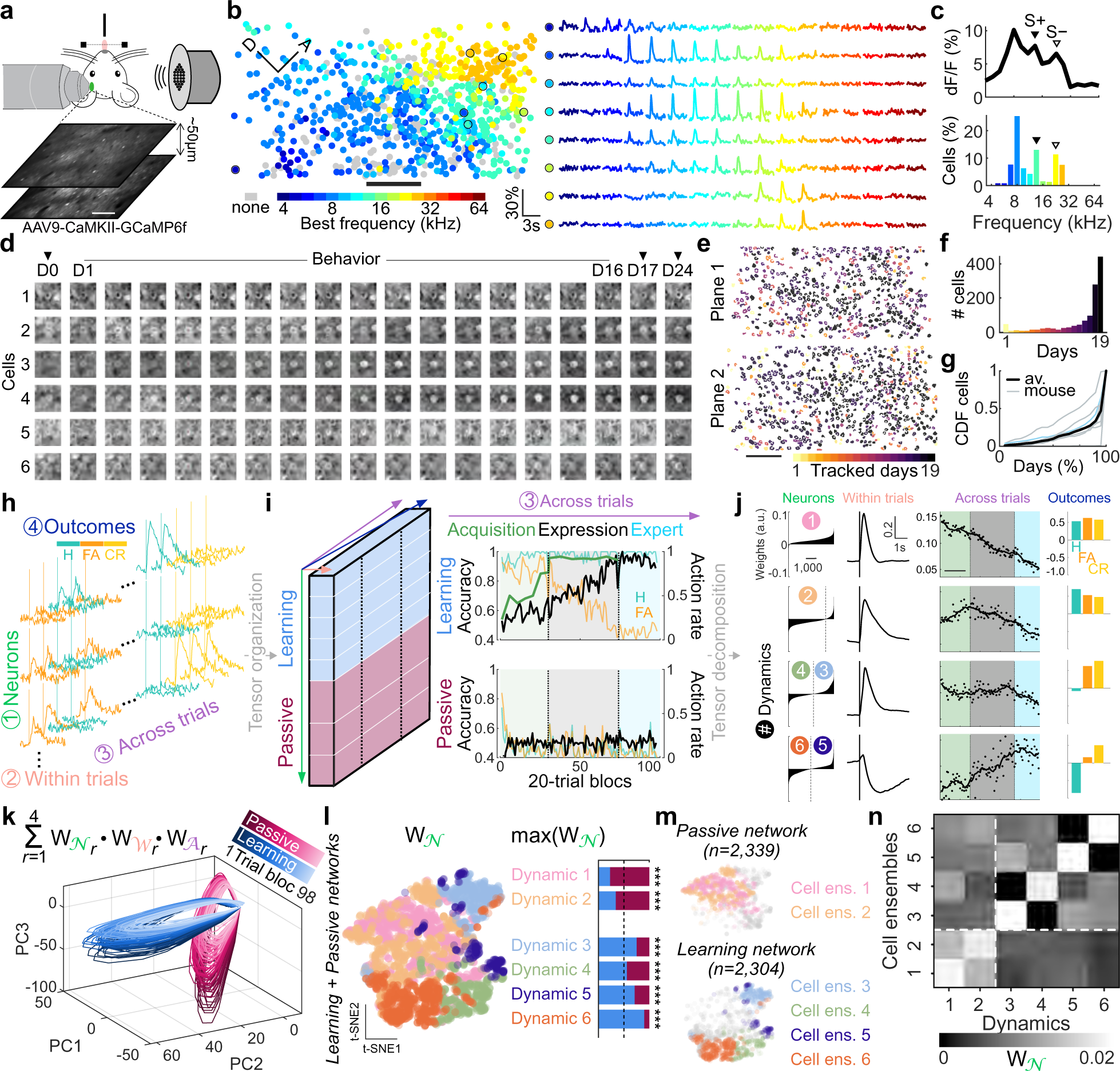

From this high-dimensional dataset, we sought to identify single neurons and neuronal ensembles carrying learning-related information, resolve stimulus and non-stimulus related activity within a given trial, identify changes in representation across trials, and determine outcome-specific dynamics. To do so, we organized our data into a 4-dimensional array containing neurons *×* time in trial *×* trials across learning *×* trial outcome (Fig. 2h). To identify shared and distinct variability in neuronal populations recorded in passive mice (*n* = 2, 339, ‘passive network’) and in learning mice (*n* = 2, 304, ‘learning network’), we created a ‘megamouse’ by combining data from all mice and aligning neural activity to learning phase (n=4,643 neurons, see Methods; Fig. 2i; Supplementary Figure 8). We then used low-rank tensor decomposition to allow unsupervised identification of demixed, low-dimensional neural dynamics across multiple (*>* 2) dimensions^26,27^ (Supplementary Figure 9 and Supplementary Figure 10a,b; see Methods). The tensor decomposition revealed six neuronal dynamics, each characterized by the four factors of the original tensor (see Methods; Figs.2j, Supplementary Figure 10c,d, Supplementary Figure 11d). These six dynamics represented independent computations performed by the auditory cortical networks.

Projecting the product of the decomposition into principal component subspace showed that learning and passive networks exhibit almost orthogonal dynamics (Fig. 2k; Supplementary Figure 10f,g) and that the neural dynamics of different trial types evolved further apart in the learning network than in the passive network (Supplementary Figure 10h,i). Importantly, we ensured that the identified dynamics were not driven by isolated mice (Supplementary Figure 10e). Therefore, decomposition of the megamouse tensor discovered distinct dynamics exhibited by passive versus learning networks.

For further analyses, we attributed each dynamic to individual neurons based on the neuron’s maximum weight (‘unique participation’; Fig. 2l; see Methods and Supplementary Figure 11). This allowed us to map the six dynamics onto six distinct cell ensembles, i.e. groups of neurons maximally encoding a particular network-specific dynamic (Fig. 2m and Supplementary Figure 11d). It is important to note that individual neurons (and corresponding ensembles) could exhibit mixed selectivity for the six dynamics, which allows an individual neurons to contribute to multiple, independent computations (Fig. 2n).

**Learning counteracts tone-evoked habituation by maintaining stimulus selectivity in distinct cell populations:** A prevailing view in sensory systems holds that sensory cortices subserve associative learning through plasticity of the cue representation^5,28–36^. This model posits that individual neurons (via changes in sensory tuning) and neural populations (via cortical map expansion) enhance the representation of behaviorally relevant cues for use by downstream regions^37–39^. These studies, however, measure neural tuning and map expansion outside of the task context in a ‘pre’ and ‘post’ learning design and infer that plasticity of cue representations reflects the mechanistic role of the sensory cortex. To assess this model, we initially focused on the cell ensembles that exhibited classical stimulus-evoked activity (Fig. 2j), namely cell ensembles 1-4.

We observed a prominent signature of stimulus-evoked habituation over hundreds to thousands of trials. This habituation dominated activity in passive networks, as seen in cell ensembles 1 and 2 which represented *∼*77% (1, 803*/*2, 339) of all passive cells (Fig. 3a,d). These neurons exhibited stimulus-evoked activation (cell ensemble 1) or suppression (cell ensemble 2), both of which decreased in amplitude over time (Fig. 3b-c,e-f). These cell ensembles were not stimulus selective and displayed the same dynamic in both stimulus 1 (S1) and stimulus 2 (S2) trials (Fig. 3b,e). These ensembles thus reflected the broad-based suppression of non-selective neurons after long-term repeated presentation of the same sounds.

Fig. 3.
Learning counteracts tone-evoked habituation by maintaining stimulus selectivity in distinct populations.
**a**, Representation of cell ensemble 1 in the Passive network. **b**, Average activity of cell ensemble 1 in S1 (black) and S2 (gray) trials across time in 80-trial blocks. Black triangles indicate tone onset, gray lines delimit averaged trial blocks. Black dashed lines separate time phases indicated by light to dark gray rectangles at the top: early, middle and late (see Methods). **c**, Cell ensemble 1 tone-evoked calcium responses across time phases for S1 and S2 trials combined (Friedman test, *p* = 1.26.10*^−^*^291^). **d**, Representation of cell ensemble 2 in the Passive network. **e**, Average activity of cell ensemble 2 in S1 and S2 trials across time. **f**, Cell ensemble 2 tone-evoked calcium responses across time phases for S1 and S2 trials combined (Friedman test, *p* = 7.32.10*^−^*^1^^21^). **g**, Representation of cell ensemble 3 in the Learning network. **h**, Average activity of cell ensemble 3 in hit (green) and CR (yellow) trials across learning in 80-trial blocks. Black triangles indicate tone onset, gray lines delimit averaged trial blocks. Black dashed lines separate learning phases indicated by colored rectangles at the top: Acquisition, Expression and Expert (see Methods). **i**, Representation of cell ensemble 4 in the Learning network. **j**, Average activity of cell ensemble 4 in hit and CR trials across learning. **k**, Response index (response probability over learning; see Methods) of cell ensembles 1 and 2 (red) vs cell ensembles 3 and 4 (blue) (Wilcoxon test, *p* = 1.23.10*^−^*^30^). **l**, Selectivity index (see Methods) of cell ensembles 1 and 2 (red) vs cell ensembles 3 and 4 (blue) (Wilcoxon test, *p* = 1.37.10*^−^*^94^). **m**, Pre (top raw) and post (bottom raw) learning tonotopic maps (left), after spatial binning (middle) and restricted to surface with S+ (filled triangle) and S*−* (open triangle) best frequency (right) of one example mouse. **n**, Change in surface representation of S+ and S*−* pre- vs post-task learning (Learning) or pre- vs post-passive exposure (Passive) (binomial proportion tests). **o**, Pre vs post-learning change in percentage of neurons responsive to S+ and S*−*(binomial proportion tests). **p**, Pre vs post-learning change in tone-evoked responses of pre-task S+ and S*−* responsive neurons (KW test, *p* = 2.77.10*^−^*^5^). **q**, Pre- vs post-learning comparison of local best frequency differences in tonotopic maps. **r**, Distribution of local differences (from difference maps in **q**) in Learning versus Passive. median *±* s.e.median; \**p <* 0.05; \*\**p <* 0.01; \*\*\**p <* 0.001, n.s.: not significant.

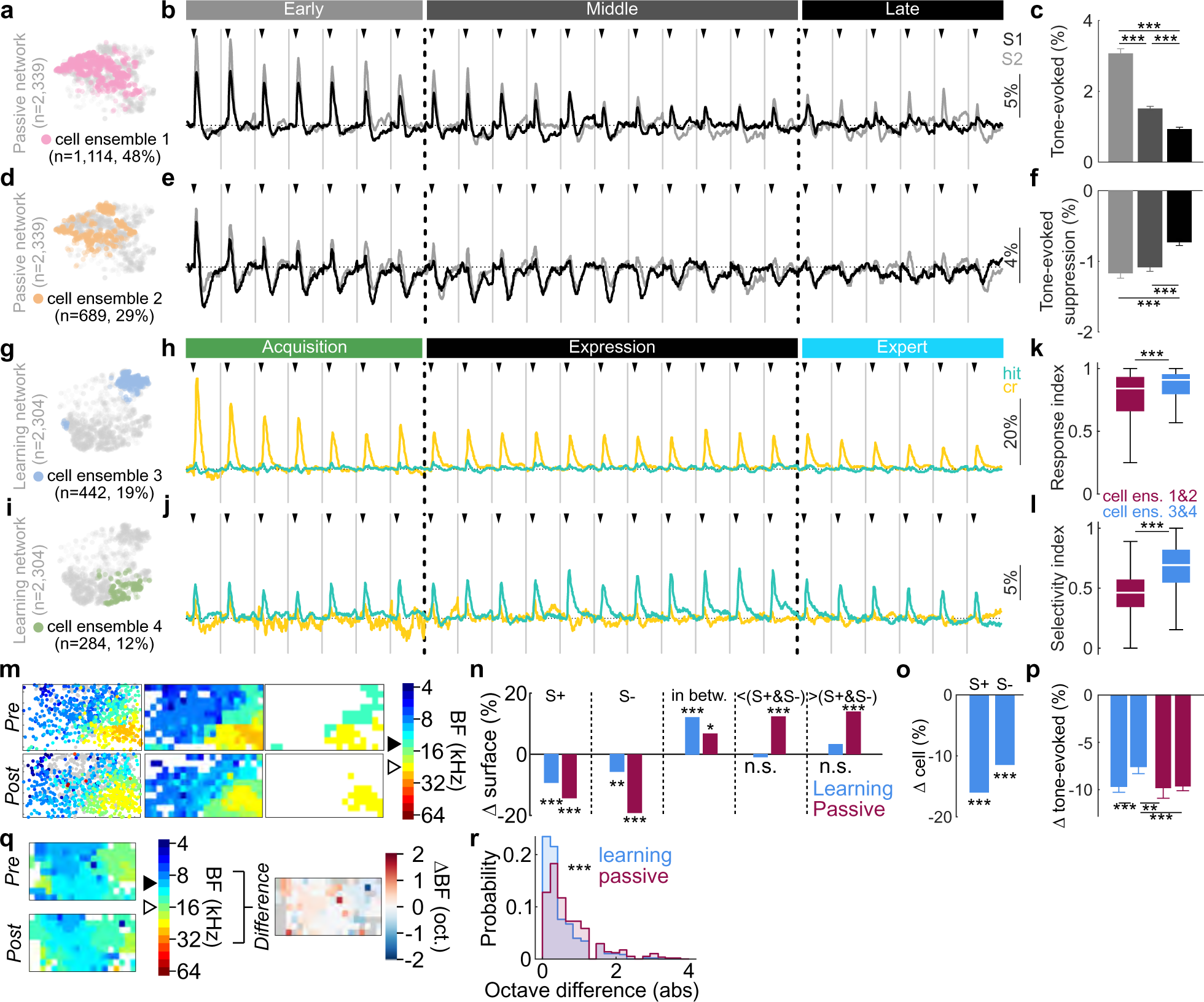

Stimulus-evoked responses in learning networks were observed in cell ensembles 3 and 4 (Fig. 3g-j). This includes a high selectivity for the S*−* (cell ensemble 3) or S+ (cell ensemble 4) cues (Fig. 3g-j). Cell ensemble 3 consisted of 19% of the Learning cell population (Fig. 3g), and displayed a slight habituation but mainly a strong preference for the S*−* throughout learning (Fig. 3h), while cell ensemble 4 (12% of total learning cells; Fig. 3j) exhibited S+ selectivity throughout learning (Fig. 3j). Cell ensembles 3 and 4 were more tone responsive and tone selective than cell ensembles 1 and 2 (Fig. 3k,l). Stimulus-evoked activity analyses across days of all recorded neurons (*n* = 7, 137) also support these results (Supplementary Figure 12, Supplementary Figure 13). Therefore, learning counteracted tone-evoked habituation by maintaining distinct ensembles that encoded either the S+ or S*−* selectively.

**Learning was not associated with cortical map expansion:** To directly test representational expansion and tuning shifts, we conducted a series of analyses focusing on stimulus-evoked responses before (pre-task) and after (post-task) learning, akin to classical measures of tuning and tonotopy. We computed the change in surface area occupied by S+ and S*−* preferring cells in tuning curve sessions, outside the task (Fig. 3m). Surprisingly, we observed no increase in the map-level representation of the S+ or S*−* after learning, and instead, observed a modest decrease (Fig. 3m-n). In addition to the best frequency representation, the fraction of neurons responding to the S+ and S*−* decreased (Fig. 3o) and the response amplitude of neurons that were initially tuned to the S+ and S*−* was lower after learning (Fig. 3p). Interestingly, while we observed no increase in representation to the S+ and S*−*, learning networks favored the representation of frequencies in between S+ and S*−*, but not higher or lower as seen in passive networks (Fig. 3n). Finally, using our passive networks as a base-case comparison, we calculated the local changes in the tonotopic map structure (Fig. 3q). Learning networks were surprisingly stable and exhibited less local changes than passive networks (Fig. 3r). These pre- vs post-learning changes in responsiveness and tonotopy thus mirrored the responsiveness observed online during learning (in dynamics 1 and 2) in a stable, tracked network (n=4,643 neurons, Fig. 3a-l), as well as when we include all neurons from each session (n=7,137 neurons) (Supplementary Figure 13). Altogether, our results suggest that cortical map expansion and changes in single-neuron tuning are unlikely to be the substrate for associative learning^40,41^.

**Tone-restricted silencing only partially impairs learning and performance:** We next sought to understand the extent to which the maintenance of stimulus-selectivity by learning networks was important to learning and performing the task. We performed daily bilateral silencing of AC during stimulus presentation throughout learning (Supplementary Figure 14a). Tone-restricted AC silencing impaired task performance throughout learning (Supplementary Figure 14b-e), task acquisition (Supplementary Figure 14f-i), and online performance during learning, with gradual fading of the effect at expert performance (Supplementary Figure 14n-q). Accuracy and action rate were not affected in reinforced light-off trials (Supplementary Figure 14j-k), but PV-ChR2 mice lick more and faster to the S*−* (Supplementary Figure 14l-m), suggesting that tone-restricted AC silencing also impaired expression, but to a lesser extent than full-trial silencing. Altogether, these results showed that information carried by the AC network in the tone-evoked window is used during learning. Interestingly, tone-restricted silencing impacted learning less than full trial silencing across nearly all measures (Fig. 1, Supplementary Figure 14), suggesting that activity *after* the tone-evoked window was critical for rapid contingency acquisition and performance during learning.

**Rapid emergence of reward prediction activity in the auditory cortex:** The sensory cortex is widely considered to be specialized for perception by interpreting complex sensory objects^42,43^ or adjusting representations of behaviorally-relevant stimuli^2,33,37,44,45^. Recent evidence, however, suggests that sensory cortical neurons directly encode non-sensory variables such as movement^46–49^, reward timing^50–53^, expectation^54,55^, and context^23,45,56–63^. Conjoint representations of sensory and non-sensory variables in the same network could further hone perception or, alternatively, subserve more integrative associative processes.

Inspection of the within-trial dynamics of learning-driven cell ensembles 5 and 6 suggested that these neurons exhibited non-canonical activity in the form of a signal that occurred late in the trial, delayed from the tone-evoked response (Fig. 2j). This late-in-trial signal increased over learning and was trial type selective (Fig. 2j). We next sought to further explore the encoding properties of these two cell ensembles. Cell ensemble 5 (*n* = 155 cells from the learning network), exhibited late-in-trial activity on hit trials (licking to the S+) that increased with learning (Fig. 4a). This delayed activity was not apparent on correct S*−*trials (correct reject, CR), where neurons exhibited classical stimulus-evoked response that habituated over learning (Fig. 4b).

Fig. 4.
Rapid emergence of reward prediction encoding drives learning.
**a**, Heat map of cell ensemble 5 activity (*n* = 155 cells) across learning phases (delimited by horizontal white dashed lines) in hit trials (20-trial blocks). White trace represents the average trial trace. Inserts (right) show average activity at time indicated by black triangles. Colored rectangles indicate learning phases: Acquisition (green), Expression (black) and Expert (blue). **b**, Heat map of cell ensemble 5 activity across learning phases (delimited by horizontal white dashed lines) in CR trials (20-trial blocks). **c**, Heat map of the activity of a fraction of cells from cell ensemble 5 (*n* = 20 cells) from one example mouse across consecutive S+ trials. Black dots indicate licks. Trial outcome is represented on the right (green circle: hit; blue stars: miss). **d**, Cell ensemble 5 activity in hit vs miss trials (time and number matched, see Methods and Supplementary Figure 15a). **e**, Area under the curve (AUC) quantification of data in gray rectangle in **d** (Wilcoxon signed rank test, *p* = 6.78.10*^−^*^21^). **f**, Procedure of reinforced and probe hit trial (H) matching. **g**, Average cell ensemble 5 activity in reinforced hit trials immediately before (black) or after (gray) probe hit trials (green). **h**, AUC quantification of data in **h** (Friedman test, *p* = 0.3071). **i**, Lick PSTHs in reinforced hit trials immediately before (black) or after (gray) probe hit trials (green). **j**, Quantification of number of licks in 1-s window post-tone (KW test, *p* = 3.18.10*^−^*^56^). **k**, Average activity of cell ensemble 5 over the first five blocks of 40-reinforced hit trials in learning. **l**, Late peak activity in HIT trials across learning phases of cell ensemble 5 (green) and low weighted cells (null, black). **m**, Procedure of reinforced and probe FA trial (fa) matching (top) and corresponding local accuracy quantification (bottom; see Methods; repeated measures ANOVA, *p* = 3.16.10*^−^*^4^). **n**, Average cell ensemble 5 activity in FA trials in the probe, non-reinforced context (orange). AUC late-in-trial (gray rectangle) compared to zero (Wilcoxon signed rank test, *p* = 1.46.10*^−^*^8^). **o**, Average activity of cell ensemble 5 (*n* = 51 cells) from one example mouse in FA trials in the reinforced context (*n* = 423) after classification based on the detection of a reward prediction signal. Bottom, average activity of FA trials with (RP+, *n* = 101) or without (RP-, *n* = 322) reward prediction signal, and activity during FA trials in the probe context (*n* = 19 trials, orange) reflecting ‘knowledge’ errors (see also Supplementary Figure 16). **p**, Heat map of the activity of a fraction of cells from cell ensemble 5 (*n* = 51 cells) from the same example mouse in **o** across consecutive FA trials in the reinforced context. Identification of a RP signal is represented by a black dot (right). **q**, Distribution of RP+ and RP*−* FA trials over learning in learning mice (binomial proportion tests, Acquisition, *p* = 1.65.10*^−^*^7^, Expression, *p* = 3.32.10*^−^*^10^, Expert, *p* = 0.22). **r**, Trial-specific closed-loop optogenetic AC inactivation over learning. **s**, Performance index (left, see Methods; two-way ANOVA, *p* = 2.11.10*^−^*^21^) and hit lick latency (right; two-way ANOVA, *p* = 0.013) in probe context in post-hit silencing experiments. **t**, Performance index (left, see Methods; two-way ANOVA, *p* = 6.36.10*^−^*^5^) and hit lick latency (right; two-way ANOVA, *p* = 0.008) in probe context in post-FA silencing experiments.

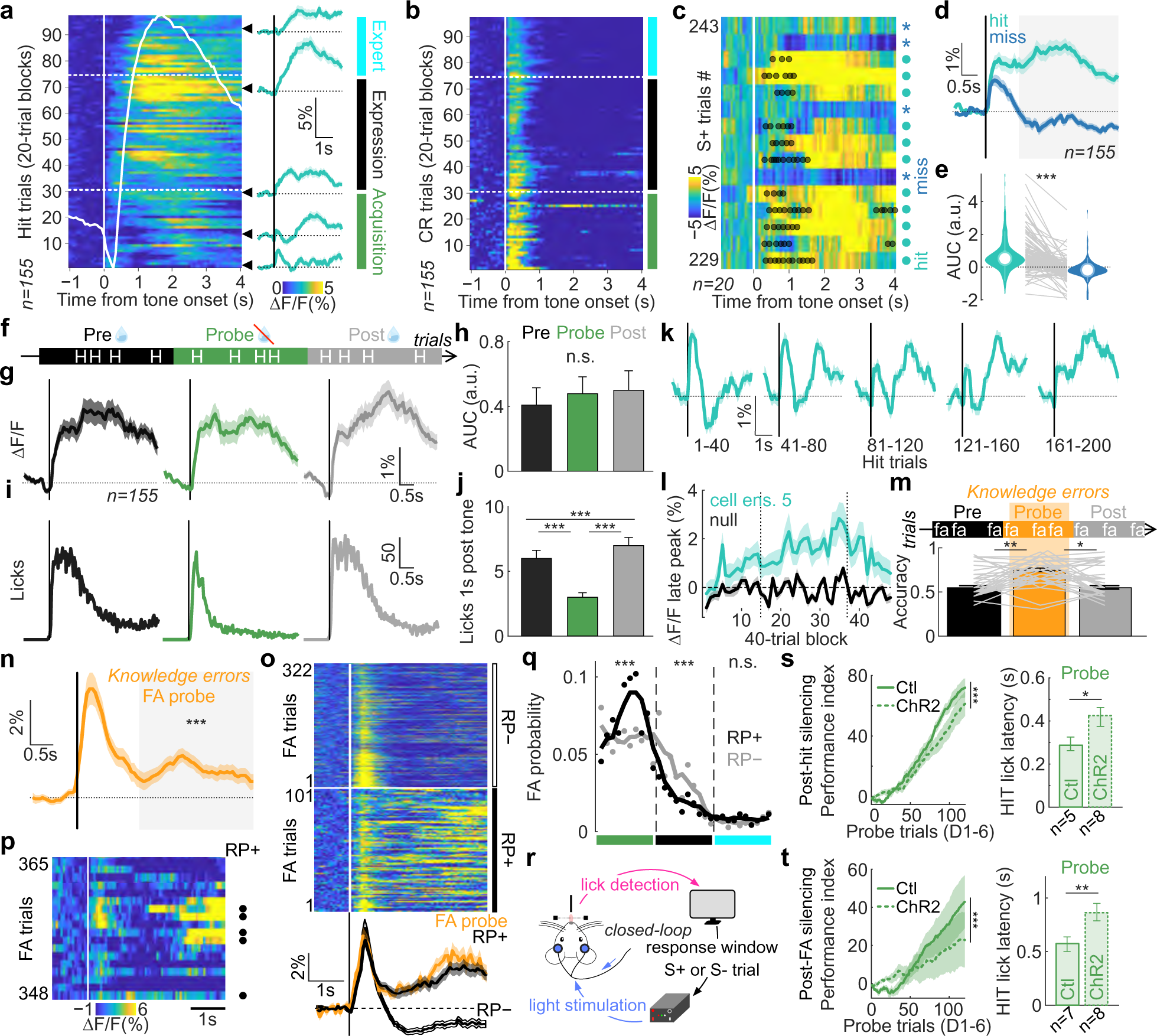

To understand the nature of the late-in-trial activity, we exploited our multiple trial types to disambiguate the contribution of sensory, motor, and reward signals. To assess whether the late-in-trial signal was a delayed form of sensory activity, we compared activity in hit trials to activity in trials where the same stimulus was presented but the mice did not lick and did not get rewarded (miss trials, Figs.1a and 4c-e). To ensure an appropriate comparison between hit and miss trials, we generated a balanced set of trials that were matched in number (given that miss trials were less frequent) and occurred within the same time period (given that the signal amplitude evolved with learning) (Supplementary Figure 15a). Cell ensemble 5 did not exhibit late-in-trial activity on miss trials (Fig. 4c-e), discarding the possibility that it reflected a delayed sensory response. We then asked whether this activity reflected reward consumption. We compared cell ensemble activity during hit trials in the reinforced context to the activity during hit trials in the probe context (Fig. 4f), where the mice expected reward and thus correctly licked to the S+ but the reward was omitted (Fig. 1b). We matched the number of trials between reinforced and probe contexts and controlled for within-session and across-session changes by comparing probe hit trials to reinforced hit trials immediately before and after the probe block (Fig. 4f). Strikingly, late-in-trial activity was preserved in probe trials (Fig. 4g,h), indicating that it did not reflect reward consumption. Finally, although movement has been reported to decrease auditory cortical activity^46,64–66^, we sought to understand the degree to which this late-in-trial signal could be driven by licking itself. To do this, we first exploited probe hit trials where the lick rate was strongly reduced compared to reinforced hit trials (Fig. 4i,j). We observed no difference in the late-in-trial neural signal and could thus conclude that the signal was not due to ongoing licking (Fig. 4i,j). Second, we tested the possibility that this late-in-trial signal was driven by the initiation of a lick bout as compared to the ongoing licking activity. We isolated spontaneous lick bouts in between training blocks and observed that the cell ensemble was not lick-responsive (Supplementary Figure 15b,c). In addition, if lick initiation drove this activity, we would also expect to see it on false alarm trials (incorrect licking to the S*−*). For this analysis, we focused on false alarms that occurred after task acquisition, as these errors are unlikely to be errors due to imperfect task knowledge. We observed no systematic late-in-trial activity on these trials (Supplementary Figure 15d) even though the licking pattern in false alarm trials was similar to that during probe hit trials (Supplementary Figure 15e). Taken together, the late-in-trial activity did not reflect stimulus, reward consumption, licking, nor lick initiation. Instead, these results showed that cell ensemble 5 encoded the higher-order process of reward prediction (RP).

We next sought to identify the precise moment when a contingency is formed by identifying the trials when this reward prediction signal emerged. Initially, these neurons exhibited classical tone-evoked responses but then abruptly and within only 40 hit trials, developed a robust reward prediction activity (Fig. 4k, Supplementary Figure 15f). This reward prediction signal continued to develop over Acquisition, strengthened during Expression, and then surprisingly receded at Expert level when learning is nominally complete (Fig. 4a,l, Supplementary Figure 15g). This longitudinal temporal dynamic mirrored our optogenetic results which demonstrates that the AC is the default pathway for learning but then becomes dispensable at expert levels. Altogether, these results show that a reward prediction signal rapidly emerges at the timescale of Acquisition in auditory cortical networks.

**Revealing the underlying cognitive drivers of errors:** Identifying the cognitive drivers of errors is particularly challenging during learning ^4^. Errors during learning are typically considered ‘mistakes’ while discriminative contingencies (task knowledge) are still forming. However, errors arise not only from knowledge-related mistakes (for which animals incorrectly expect reward), but also from factors such as impulsivity, disengagement, and exploration (for which animals do not expect reward). While detailed behavioral inspection has been a promising route to uncover the nature of errors^11^, an alternative approach is to use neural activity itself. Given our findings of reward prediction encoding on correct trials, we hypothesized that the same signal would be present when animals make ‘knowledge-related’ errors, when animals incorrectly ‘expected’ rewards on S*−* trials. To address this, we first focused on the occasional false alarms (FA) that occurred during probe trials, as they reflected errors of task knowledge (Fig. 4m)^7^. Strikingly, we observed a robust reward prediction activity in these trials (Fig. 4n), strongly suggesting that animals were indeed expecting reward. We next reasoned that such knowledge errors should be present not only on probe trials, but also in a subset of reinforced trials, interspersed with non-knowledge errors. We classified individual FA trials in the reinforced context based on the presence of a reward prediction signal (see Methods; Supplementary Figure 16a). We identified a significant proportion of trials that exhibited robust reward prediction activity, but also many that did not (Fig. 4o, Supplementary Figure 16b). The reward prediction signal was identical to that observed in probe trials (Fig. 4o, Supplementary Figure 16d), providing further confidence that these were indeed knowledge errors. These data suggest that we could isolate knowledge errors using neural data, which was not possible from behavioral inspection alone (Supplementary Figure 16c). Interestingly, we found that knowledge errors were interspersed with errors that did not elicit reward prediction activity (Fig. 4p). Finally, we hypothesized that knowledge errors should predominantly occur during the Acquisition phase of behavior, when animals are still learning the discriminative contingencies. We computed the fraction of RP+ (knowledge-related errors) and RP-(non-knowledge errors) over time and found that RP+ errors peaked during the Acquisition phase of learning, and rarely occurred during Expression or Expert phases of behavior (Fig. 4q, Supplementary Figure 15d). These results demonstrate that the internal cognitive drivers of errors may be accessible from neural data, which is particularly striking when behavior alone is insufficient.

**Reward prediction activity provides the core teaching signal:** Learning theory proposes that animals learn from correct actions that are rewarded but also from incorrect actions that are not rewarded^67^. This allows animals to select the appropriate action after reward-predictive (S+) versus non-predictive (S*−*) cues. Given the presence of the reward prediction activity on correct S+ trials (throughout learning) and incorrect S*−* trials (early in learning), we reasoned that silencing auditory cortical activity during the post-response period could impact learning and/or performance. To test this, we performed closed-loop probabilistic optogenetic silencing of the AC whereby light was delivered upon lick detection in 90% of either S+ reinforced trials (*n* = 5 control, *n* = 8 PV-ChR2 mice) or, in a separate cohort, S*−* reinforced trials (*n* = 7 control, *n* = 8 PV-ChR2 mice; see Methods; Fig. 4r, Supplementary Figure 17a, Supplementary Figure 18a). No light was delivered in 10% of S+ reinforced trials and 100% of probe trials. Given that the light was delivered after the instrumental lick response, the effect of the manipulation could not affect the instrumental behavior on the current trial, only on subsequent ones. To confirm this, we calculated the difference in performance between light-on and light-off trials and observed no difference (Supplementary Figure 17b-d and Supplementary Figure 18b-d). In the S+ cohort, post-hit silencing weakened the stimulus-action association (Fig. 4s), delayed cue-response discrimination (Figs.4s), but did not impact probe accuracy over the first 6 days (Supplementary Figure 17e-g). Importantly, the same silencing protocol above the visual cortex (*n* = 6 PV-ChR2 mice) had no effect on behavior, confirming that these effects were specific to the auditory cortex (Supplementary Figure 17k,l). In the S*−* cohort, post-FA silencing weakened the stimulus-action association as measured on hit trials (Fig. 4t), robustly delayed cue-response discrimination (Fig. 4t, Supplementary Figure 17g), and impaired probe accuracy over the first 6 days (Supplementary Figure 17e,f). Accuracy of PV-ChR2 mice was lower than control in the reinforced context in both experiments (Supplementary Figure 17h and Supplementary Figure 18h), with lower hit rate and higher FA rate (Supplementary Figure 17i and Supplementary Figure 18i), and longer response latencies on hit trials (Supplementary Figure 17j and Supplementary Figure 18j), suggesting an impairment of expression. Overall, these closed-loop manipulations showed that AC activity at the time of the reward prediction signal in both hit and FA trials was used by the animal for the task acquisition and expression. These data also demonstrate that learning is sensitive to cortical silencing on mistakes (FA trials) suggesting that in a go/no-go paradigm, reward feedback on error trials is crucial to the learning process. Altogether, these results suggest that reward prediction activity in auditory cortical networks is used as a teaching signal during learning.

**Encoding of action suppression enables task performance:** A critical requirement in a go/no-go task is the ability to suppress responding to the non-rewarded, S*−* cue. In our task, we demonstrate that mice have the capacity to withhold licking to the S*−* very early in learning (as shown in probe trials during the acquisition phase) but continue to lick for hundreds to thousands of trials when being reinforced and throughout Expression. Here, we ask the extent to which the AC mediates this form of action suppression. Neurons in cell ensemble 6 (*n* = 704, 31% of learning networks; Fig. 5a), but not non-member cells, exhibited late-in-trial activity when animals correctly withheld from licking on S*−* trials (correct rejects, CR; Fig. 5b, Supplementary Figure 19a-b). This signal was stable throughout training despite the strong increase of CR rate over learning (Fig. 5c, Supplementary Figure 19c-d). This all-or-none attribute suggested that this late-in-trial activation was tied to performance rather than being a signal used for learning. Once mice acquired the task contingencies, they essentially learned to inhibit a licking response to the S*−* tone. We therefore thought to test the hypothesis that late-in-trial activation in CR trials reflected action suppression. First, we reasoned that activity in FA and CR trials should be similar until the moment of suppression failure (i.e. first lick). We compared the activity of cell ensemble 6 in CR vs FA trials, i.e. when mice fail to withhold licking (see Methods) exploiting the different first lick latencies in FA trials (Fig. 5d). We observed that calcium activity dropped abruptly in FA trials at the time of the first lick compared to CR trials (Fig. 5d,e, Supplementary Figure 19e). Second, if lick suppression is an active contingency-specific process, the late-in-trial activation should be specific for correct rejections for the S*−* tone, and not observed when the animal did not lick in response to the S+ tone (miss trials). Given that miss trials were rare and sporadic, we controlled for the effect of time over learning and difference in the number of trials for each outcome type (see Methods) and did not observe late-in-trial activation on miss trials despite similar peak activity after tone onset in miss and CR trials (Fig. 5f). Third, we reasoned that if this activity reflects the active process of action suppression, the signal should decrease when the animal disengaged from the task. We therefore compared late-in-trial activity in CR trials immediately before, during and after short blocks of disengagement (see Methods) and observed that the activity dropped significantly when mice transiently disengaged from the task (Fig. 5g). These data suggest that the auditory cortex integrates a higher-order action suppression signal.

Fig. 5.
Action suppression signals in the AC induce suppression of licking.
**a**, Representation of cell ensemble 6 (*n* = 704 cells) in the Learning network. **b**, Average activity of cell ensemble 6 (yellow) versus cells that do not contribute to this dynamic (null, black) in CR trials in Expert phase (Wilcoxon test, *p* = 7.44.10*^−^*^17^). **c**, Average activity of cell ensemble 6 in CR trials (top) and CR rate (bottom) during Acquisition (green), Expression (black) and Expert (blue) phases (KW test, *p* = 0.09). **d**, Heat map of cell ensemble 6 activity in hit, FA and CR trials. FA trials are binned according to lick latencies (white dots, latency range extrema; white cross, mean latency). **e**, Heat map of cell ensemble 6 activity in hit and FA trials significantly different from CR trials (Wilcoxon tests, red, higher; blue, lower; white, n.s.). **f**, Average cell ensemble 6 activity in miss and CR trials (time and number matched, see Methods; middle). Quantifications of tone-evoked activity (bottom left; Wilcoxon signed rank test, *p* = 0.84) and late-in-trial AUC (bottom right; Wilcoxon signed rank test, *p* = 5.24.10*^−^*^26^). **g**, Procedure of reinforced and probe CR trial matching (top) and corresponding calcium activity (middle; Friedman test, *p* = 1.36.10*^−^*^11^) and local hit rate (bottom; Friedman test, *p* = 3.45.10*^−^*^11^). **h**, FA rate difference between light-on and light-off trials in PV-ChR2 mice (two-way ANOVA, *p* = 7.20.10*^−^*^16^; t-tests compared to 0, *p* = 4.96.10*^−^*^4^, *p* = 0.96, *p* = 0.002). Auditory or visual cortex were inhibited during the full trial (AC trial, *n* = 8; VC trial, *n* = 8) or AC was silenced during tone presentation only (AC tone, *n* = 4). **i**, Average lick probability in FA light-on versus FA light-off trials (two-way ANOVA, *p* = 1.18.10*^−^*^5^; t-tests compared to 0, *p* = 1.94.10*^−^*^6^, *p* = 0.10, *p* = 0.68).

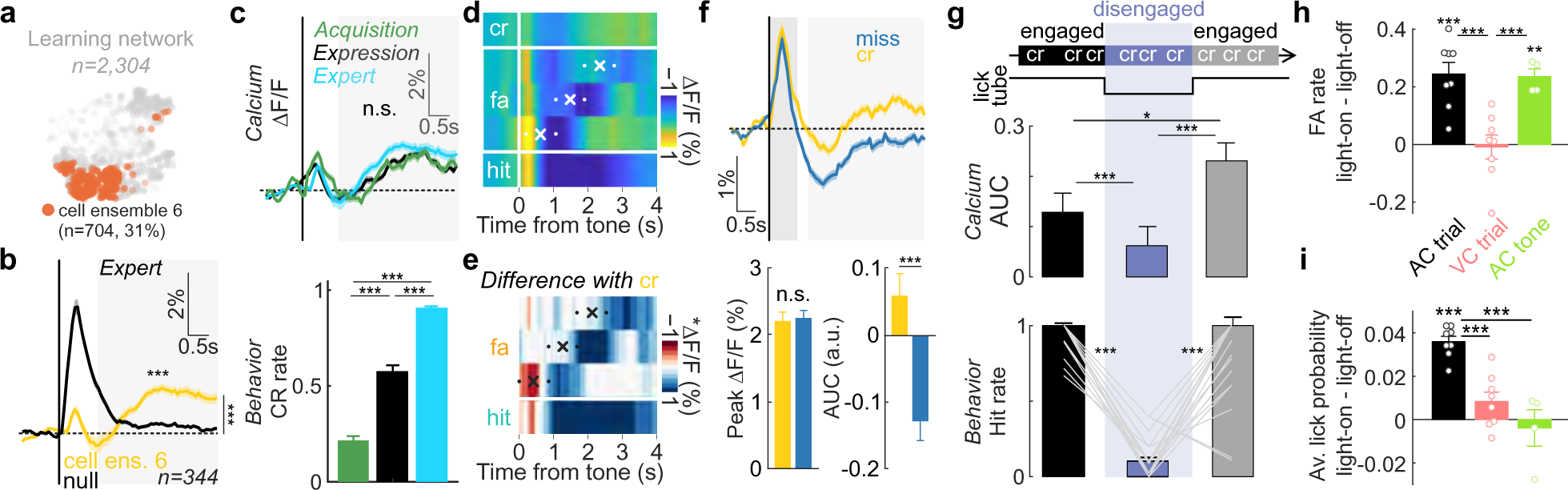

Finally, we wondered whether the action suppression activity in the AC was causal to performance during learning. To test this, we reasoned that silencing the AC network throughout S*−* trials should increase the FA rate but also the lick probability (since the action suppression neurons are silenced during this period). In contrast, silencing the AC network only during the stimulus period should increase the FA rate but not impact the lick probability when the light is off (Supplementary Figure 19f). We first compared the FA rate between light-on and light-off trials in PV-ChR2 mice during full trial silencing (Fig. 5h) and observed a marked increase in FA rate and lick probability (Fig. 5h,i). Importantly, this effect was not the result of the perception of optogenetic manipulation *per se* as suppression of the visual

**Higher-order contingency ensembles are spatially clustered and uncoupled from sensory representations:** We next asked the extent to which reward prediction and action suppression ensembles mapped onto the underlying stimulus properties of the AC. We exploited the spatial resolution of two-photon imaging to characterize the spatial distribution of reward prediction and action suppression neurons in the AC network. Strikingly, we observed that the two cell ensembles were spatially clustered (Fig. 6a-c). To determine whether this organization was driven by the neuron’s pre-learning stimulus selectivity, we calculated the selectivity index (SI) of each neuron before training to test whether neurons selective for the S+ preferentially became reward prediction neurons and S*−* selective neurons preferentially became action suppression neurons. We observed no difference in SI distribution between reward predictive and action suppression neurons (Fig. 6d,e), suggesting that pre-task stimulus selectivity was not predictive of either reward prediction or action suppression. We then asked whether the spatial location of reward prediction and action suppression neurons aligned with the underlying tonotopic map. In other words, did action suppression neurons have S*−* tone for best frequency, and were reward prediction neurons preferentially responsive to S+ tone (Fig. 6f; see Methods)? We found that this was not the case (Fig. 6g,h), with similar proportion of S+- and S*−*-preferring neurons in reward prediction and action suppression cell ensembles (Fig. 6i). Therefore, contingency-related ensembles clustered into spatial domains that were uncoupled from underlying stimulus selectivity and tonotopy, indicating a higher-order functional segregation within the AC.

Fig. 6.
Reward prediction and action suppression signals emerged in segregated neuronal populations and do not rely on underlying stimulus selectivity
**a**, Spatial distribution of reward prediction (purple circles) and behavioral inhibition (orange circles) cell ensembles in an example mouse. Color scale indicates neuronal weights in Dynamics 5 (purple) and 6 (orange). **b**, Median of cell distance between cell ensembles compared to shuffle distribution (*n* = 500) for example mouse in **a**. The null hypothesis is that the distance between the two ensembles is no different than chance (i.e. no spatial organization). **c**, Z-scored distances between clusters per mouse (blue: significant; gray: non-significant). Red arrow points to example mouse in **a**. **d**, Neuronal weights in Dynamics 5 and 6 of cells from learning mice (*n* = 1, 216, left) and their pre-task stimulus selectivity index (right). **e**, Distributions of pre-task stimulus selectivity of cell ensembles 5 and 6 (KS test, *p* = 0.25, Wilcoxon test, *p* = 0.18). **f**, Pre-task tonotopic map of the example mice in **a**. Cells are colored according to their best frequency (BF). Frequencies used as S+ and S*−* for training are indicated by full and empty triangles, respectively. **g**, Distribution of BF distance from S+ for reward prediction cell ensemble (purple). Null hypothesis is that reward prediction cells have a BF as close to S+ as possible (black; see Methods; KS test, *p* = 3.81.10*^−^*^9^). **h**, Distribution of BF distance from S*−* for action suppression cell ensemble (orange). Null hypothesis is that action suppression cells had a BF as close to S*−* as possible (black; see Methods; KS test, *p* = 9.21.10*^−^*^16^). **i**, Proportions of S+ and S*−*-preferring cells in reward prediction and action suppression cell ensembles (binomial proportion tests, S+, *p* = 0.17, S*−*, *p* = 0.53) cortex in PV-ChR2 mice did not have this effect (Fig. 5h,i). In contrast, restricting silencing to the stimulus period increased FA rate while not affecting lick probability (Fig. 5h,i), suggesting that the late-in-trial activity in CR trials was critical for the maintenance of action suppression. Altogether, these results showed that action suppression is encoded in the auditory cortex and is instrumental for performance during learning.

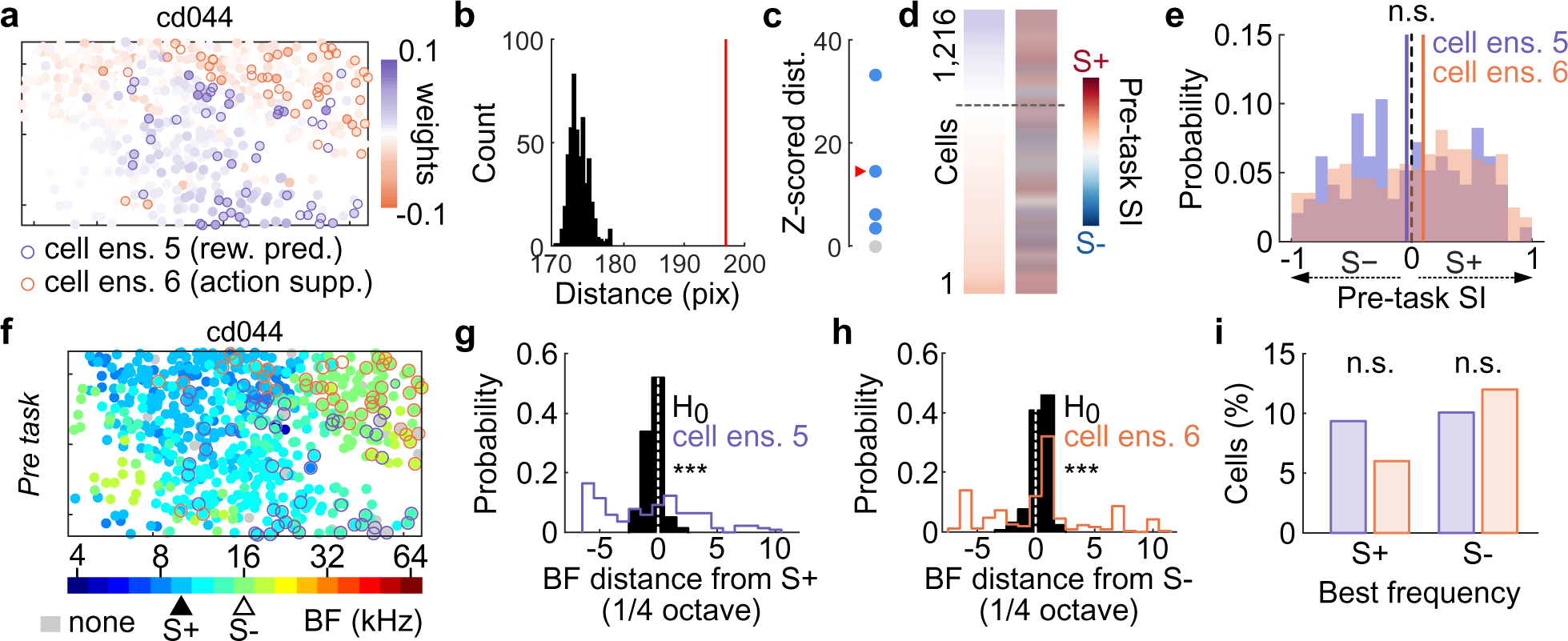

## Discussion

Learning-related neural dynamics are traditionally defined as task-specific neural activity changes that occur at the timescale of an animal’s performance improvements in the learning, i.e. a reinforced context^8^. Using this conceptual and experimental framework, perceptual and instrumental (reward-based) learning and their underlying neural dynamics have been described as slow and gradual e.g^2,44^, with animals requiring thousands of trials to learn low information-content tasks^3,4^. We took advantage of a recent behavioral paradigm^7^ that uses non-reinforced probe trials to show that task knowledge emerges more rapidly and earlier than behavioral performance improvement in the learning, reinforced context. Using this powerful behavioral manipulation to quantitatively assess when the animals acquired the task contingencies, we aligned our neuronal recordings to learning stages between animals while preserving trial-based resolution, and took advantage of an unsupervised, dimensionality reduction method across multiple timescales ^26^ to identify learning-specific neural dynamics. We observed that reward prediction activity emerged remarkably fast - within tens of trials and on the first day of training - in the AC, hundreds to thousands of trials before noticeable performance improvements. The AC thus exhibits latent knowledge of the task (encoded in the network but not behaviorally apparent) with animals experiencing periods when knowledge of environmental contingencies (between cues, actions, and rewards) becomes rapidly encoded in the brain, perhaps reflecting an insight-like moment. The latent task knowledge was manifested not as changes in sensory representations, but as the emergence of discrete ensembles encoding reward prediction (needed for identifying that a particular cue signals reward availability) and action suppression (needed for suppressing licking on S*−* trials). These computations were spatially clustered and developed in a manner that was uncoupled from the underlying stimulus-related processing that takes place in the AC, suggesting a higher-order functional organization. Overall, we find that AC contains separable and causal neural dynamics for both learning and performance.

Our results call for a revision of the classical view of the sensory cortex, according to which its primary role is to process and interpret sensory stimuli. We propose instead that the sensory cortex is better described as a sensory-enriched associative cortex, driving rapid forms of associative learning and where sensory and associative functions are intrinsically inter-mingled (i.e. co-exist within the same network) but computationally separable (Fig. 6). This function of the sensory cortex may have thus far been obscured by the use of complex sensory objects that recruit the sensory cortex for object-level processing, making it difficult to isolate non-perceptual learning computations. Finally, it is important to note that our results do not contradict studies that demonstrate single-neuron tuning curve shifts and tonotopic map plasticity when animals learn perceptually challenging tasks. Our revised model of the sensory cortex would suggest that perceptual sharpening and complex object processing can be subserved by stimulus-related plasticity while the higher-order computations related to associative learning and performance occur in parallel. We expect this view will apply beyond rodents, as rich encoding of non-sensory and task-relevant variables has also been described in human and non-human primate sensory cortical areas^68–70^.

The detailed input-output circuit that enables reward prediction and action suppression computations remains an important area for future exploration. One possibility is that ascending neuromodulatory inputs^23,29,71–75^ and top-down projections from motor and frontal regions^44,76–78^ serve as critical non-sensory inputs to the sensory cortex. The sensory cortex may then integrate and generate higher-order computations that are incorporated by broader decision-related circuits (e.g. frontal cortex, striatum and amygdala) to enable rapid learning and ongoing performance.

## Acknowledgments

We thank P. Janak, C. Honey, S. Mysore, D. Lee and the members of the Kuchibhotla laboratory for comments on the manuscript. This work was supported by the Johns Hopkins Kavli NDI Distinguished Postdoctoral Fellows Program (C.D.), National Institute of Health K99 DA059024 (C.D.), Johns Hopkins Kavli NDI Distinguished Graduate Student Fellowship (Z.Z.), National Institute of Health R00 DC015014 and R01 DC018650 (K.K.), National Science Foundation CAREER 2145247 (K.K) and Brain & Behavior Research Foundation NARSAD (K.K).

## Author contributions

C.D. and K.K. designed the experiments and data analyses; C.D. and Z.Z. performed the two-photon experiments; C.D., Z.Z. and K.F. preprocessed calcium imaging data; C.D., Z.W., K.F., A.W., and S.E. performed the optogenetic experiments; C.D. analyzed the data; C.D. and K.K. wrote the manuscript.

## Competing interests

The authors declare no competing interests.

## Materials & Correspondence

Correspondence and material requests should be addressed to.

## Methods

### Animals

All procedures were approved by Johns Hopkins University Animal Care and Use Committee (MO20A272). Male and female double (PV-ChR2; test mice) or single (PV-cre or flox-ChR2; control mice) transgenic mice between 6 and 12 weeks at the start of experiments were used for the optogenetic experiments. PV-cre (Jackson laboratory, strain #017320), flox-ChR2 (Ai32, Jackson laboratory, strain #012569) and PV-ChR2 mice were bred in-house. PV-ChR2 mice were obtained by crossing male PV-cre^+/^*^−^* mice with female flox-ChR2^+/+^ or by crossing male flox-ChR2^+/+^ with female PV-cre^+/^*^−^*. To obtain PV-cre^+/^*^−^* line, we bred female PV-cre^+/+^ with male C57BL/6J (Jackson laboratory, strain #000664). Offspring genotypes were confirmed by PCR (Lucigen EconoTaq Plus GREEN 2X) and using two-photon imaging to observe expression of the reporter protein (GFP, see subsection ‘Optogenetic experiments’). Male C57BL/6J (Jackson laboratory, strain #000664) aged between 6 and 12 weeks at the start of experiments were used for two-photon calcium imaging experiments. Animals were group housed in standard plastic cages with food available *ad libitum* and maintained on a 12-hour reversed light-dark cycle at stable temperature (19.5-22*^◦^*C) and humidity (35-38%). Experiments took place during the dark phase. Mice were kept on a mild water restriction diet (*>*85% of body weight) after surgery and throughout task training.

### Surgical procedures

Mice were anesthetized with isoflurane (5% at induction and maintained at 2% during surgery) and their body temperature was maintained at *∼*35*^◦^*C throughout the surgery.

#### Calcium imaging experiments

Mice were injected (34 gauge, 25.4 mm, 12-degree bevel needle; Hamilton Company) with 1*µ*l of AAV9-CaMKII-GCaMP6f (Addgene, #100834-AAV9, dilution 1/15) at 0.75*µ*l.min*^−^*^1^ (microinjection pump, Harvard Apparatus) in the left primary auditory cortex (centered at 1.75 mm anterior to the intersection of the lambdoid and interparietal-occipital sutures, DV:-200*µ*m). Above the injection coordinates, a cranial window was implanted replacing a circular piece of skull by a 3-mm diameter cover glass slip (Warner Instruments) that was secured in place using a mix of dental cement and Krazy Glue. A custom-made, three-point stainless steel headpost was secured to the skull with C&B Metabond dental cement (Parkell). The headpost consisted on a two-point kinematic fixation on the right side of the head, prolonged by a rod encircling the cranial window and descending at *∼*45*^◦^* ventrally on the left. Mice were given a two-week recovery period to allow weight recovery and viral expression.

#### Optogenetic experiments

3-mm diameter cover glass slips were implanted bilaterally over the auditory cortex (centered at 1.75 mm anterior to the intersection of the lambdoid and interparietal-occipital sutures, on the ridge line of the temporal bone). Custom-made aluminum funnels were implanted above each cranial window. The role of these funnels was threefold: 1) to precisely center the end of the patch cord on the cranial windows, 2) to hold the patch cord perpendicular to the cranial window (optimizing in-depth light diffusion), and 3) to fix the distance between the patch cord and the cranial window to allow identical light delivery across days. A custom-made, two-point stainless steel headpost was fixed onto the skull with C&B metabond dental cement (Parkell) and dental cement. Mice were allowed to recover for at least one week following surgery.

#### Optogenetic silencing verification experiments

For silencing verification experiments (Supplementary Figure 2, n=2), PV-ChR2 mice were injected with 1*µ*l of AAV-CaMK2-GCaMP6f (Addgene, #100834-AAV9, dilution 1/15) at 0.75*µ*l.min*^−^*^1^ in the left primary auditory cortex (centered at 1.75 mm anterior to the intersection of the lambdoid and interparietal-occipital sutures, DV:-200*µ*m) and implanted with a 3-mm cover glass slip and a custom-made, two-point stainless steel headpost. Mice were given a two-week recovery period to allow weight recovery and viral expression.

### Auditory Go/No-go task

All mice (optogenetic and two-photon imaging) underwent the same habituation and training procedures. After recovery from surgery, mice were water restricted for at least 5 days so that their weight stabilized at 85% of their *ad libitum* weight. During this period, mice were handled daily. Mice were then head-fixed and placed in the experimental context, where they were trained to lick from a lick tube or water cup to receive a drop of water (3*µ*l). No tone was presented during lick training. Lick training session ended after 30 min or when 1 ml of water was consumed. After two days of lick training, mice were trained on the auditory Go/No-go task for at least 15 days.

Mice were trained to lick to a target (S+) tone to receive a water drop (3*µ*l) and withhold licking to the foil (S*−*) tone to avoid a timeout. Auditory stimuli were three quarter octave-spaced pure tones. Target and foil tones were presented pseudo-randomly and counterbalanced every 20 trials. Each trial consisted of a no lick period (1 s), tone presentation (100 ms), dead period (200 ms), response period (2.5 s) and a delay period: hit: 4 s (to enable full licking of the reward), miss and correct reject: 2 s, false alarm: 7 s (timeout). In this learning context, called the ‘reinforced’ context, the lick-tube delivering water was positioned within reach of the tongue. In contrast, in the ‘probe context’, the lick-tube was moved out of tongue and whisker reach by an automated actuator. The blocks of probe trials were interspersed between reinforced trials and no additional delay was introduced by lick-tube movement. Importantly, we have shown that the performance gap observed between probe and reinforced trials early in learning is not driven by the change in the sensory context induced by the absence of the lick-tube in the probe context^7^.

### Optogenetic experiments

Mice were trained in the Go/No-go task for 300 trials every day: 280 trials in the rein-forced context interspersed with a short block of 20 non-reinforced (probe) trials starting at trial #141. Head-fixation habituation, lick training and Go/No-go task training took place in custom-made, sound-attenuated behavioral boxes (ambient noise level *∼*53 dB SPL) controlled with custom-written MATLAB programs interfacing with Bpod State Machines (Sanworks). Pure tones (4,757 and 8,000 Hz) were delivered through an electrostatic speaker driver (TDT) to a free field electrostatic speaker (TDT) at an intensity of 70 dB SPL and licks were detected through an infrared beam. Blue light (453nm, DPSS laser, Opto-Engine LLC) was delivered in a 20-Hz sinewave generated by Arduino. The power recorded at the end of the patch cord (splitter branching fiber-optic patch cords, Doric Lenses) was 6-8mW. When dispersed over a diameter of 3mm, that yields a light intensity of 0.85-1.13 mw/mm^2^ at the cortical surface. Sound amplitude, water drop size, and laser power were calibrated at the beginning of each experiment. To dissociate the effect of AC silencing on behavior from its consequence on the learning process, we used a probabilistic approach whereby no light was delivered during probe trials and a subset of reinforced trials. These light-off trials were critical to assess behavior when the auditory cortex was available again.

<u>Full trial experiment (</u>*<u>n</u>* <u>= 8 PV-ChR2,</u> *<u>n</u>* <u>= 8 control mice,</u> *<u>n</u>* <u>= 8 PV-ChR2 visual cortex):</u> light was turned on on 90% of reinforced trials pseudo-randomly (18 trials – 9 S+ and 9 S*−* – every 20-trial block). In light-on trials, the light was turned on 100 ms before tone onset and stayed on for *∼*2.5 s for all trial types (hit: 2.5 s post operant lick, CR and miss: stop at the end of response window, FA: 2.5 s post first lick).

<u>Expert only full trial experiment (</u>*<u>n</u>* <u>= 4 PV-ChR2 mice):</u> Mice were trained for 18 days without light. Afterward and for 5 days, from day 19 to 23, the light was turned on following the ‘full trial experiment’ protocol or on 90% of reinforced trials consecutively.

<u>Tone experiment (</u>*<u>n</u>* <u>= 4 PV-ChR2,</u> *<u>n</u>* <u>= 3 control mice):</u> light was turned on on 90% of reinforced trials pseudo-randomly (18 trials – 9 S+ and 9 S*−* – were light-on every 20-trial block). In light-on trials, the light was turned on 100 ms before tone onset and turned off at tone offset.

<u>Post hit experiment (</u>*<u>n</u>* <u>= 8 PV-ChR2,</u> *<u>n</u>* <u>= 5 control mice,</u> *<u>n</u>* <u>= 6 PV-ChR2 visual cortex):</u> we used a closed-loop lick-triggered stimulation approach, whereby light was turned on after a rewarded lick on 90% of reinforced trials pseudo-randomly (light could be turned on on 9 over 10 S+ trials every 20-trial block). In light-on trials, the light was turned on 70 ms after the first lick detection (to allow the lick cycle to complete and the tongue to retract) and 100 ms before reward delivery and stayed on for 2.5 s.

<u>Post false alarm experiment (</u>*<u>n</u>* <u>= 8 PV-ChR2,</u> *<u>n</u>* <u>= 7 control mice):</u> we used a closed-loop lick-triggered stimulation approach, whereby light was turned on after a non-rewarded lick on 90% of reinforced trials pseudo-randomly (light could be turned on on 9 over 10 S-trials every 20-trial block). In light-on trials, the light was turned on 70 ms after the first lick detection (to allow the lick cycle to complete and the tongue to retract) for 2.5 s.

At the end of the experiments, mice were anesthetized (isoflurane 5% at induction and 2% during surgery; body temperature maintained at *∼*35*^◦^*C) and the left funnel was drilled out. Mice were then put under the two-photon microscope and the field of view was excited at 980nm. Green fluorescence was detected in test mice (ChR2-EYFP) but not in control mice. This procedure allowed to confirm mice genotypes and to assess cell health. Z-stacks were collected (unidirectional, 30.98 Hz; magnification 1.7 or 2.0X; range: 450*µ*m, step: 10 *µ*m, 50 frames per step; depth from brain surface 420-445 *µ*m) to generate 3D reconstruction (ImageJ).

### Longitudinal two-photon calcium imaging during learning

Two-photon fluorescence of GCaMP6f was excited at 980nm using a mode locked Ti:Sapphire laser (Spectra-Physics) and detected in the green channel (GFP emission). Imaging was performed with a two-photon resonant-scanning microscope (Neurolabware) equipped with a water immersion objective (16x, 0.8NA, Nikon) tilted to an angle of 40-50*^◦^* to image the auditory cortex. The arm of the microscope was enclosed in a custom-made sound-attenuated box. An electronically tunable lens was used to record near-simultaneously two planes in layer 2/3 (150-250*µ*m below dura, 50*µ*m spaced, 312×192*µ*m^2^, at 15.96Hz per plane, with a laser power of *≤*40 mW). Images were collected at 1.7x or 2x magnification using ScanBox (Neurolabware) and task events (sounds, rewards, licks and frames) were recorded using a digitizer (Digidata 1550b). Pure tones were delivered through an electrostatic speaker driver (RZ6, TDT) to a free field electrostatic speaker (TDT) located at *∼*5cm from the right ear at intensity of 70dB SPL. Licks were detected through an infrared beam. Scanner noise (8kHz) was attenuated using a custom-made foam sound enclosure directly surrounding the animal and the resonant scanner was set to continuous throughout the recording session (to avoid any scanning onset-related activity). Custom-written MATLAB program interfaced with RPvdsEx to control task events. Mice were placed in a plastic tube and head-fixed via a two-point pneumatic clamp on the right and a one-point, 360*^◦^*-rotational clamp on the left (at 45-50*^◦^* in the horizontal plane). The whole behavioral platform was installed on a rotation platform so that the field of views could be precisely retrieved one day to the next. Imaging fields were retrieved every day before task training by visual inspection (see also ‘Pre- and post-task tonotopic mappings’). Typically, mice were trained for three blocks of 80-100 trials, with either two blocks of 10 probe trials interleaved in two of these three blocks, or one block of 20 probe trials. The field of view was adjusted in between blocks to compensate for z-drift, if necessary. An additional 10,000 frames of spontaneous activity were recorded in a separate block at the end of each behavior session.

### Pre- and post- task tonotopic mapping

One day before lick training, mice were placed under the microscope and were presented with a set of 17 pure tones (duration 100ms), three-quarter octave spaced, in a pseudo-random order ranging from 4 to 64 kHz at 70 dB SPL. Target and foil tones were selected for the Go/No-go task as pure tones that were similarly represented in the recorded neuronal population. The same mapping procedure took place immediately after or one day after the last behavior session, and 7 and 14 days later.

### Two-photon calcium imaging and one-photon blue light stimulation for silencing verification

To validate our optogenetic silencing protocol and determine light power to use for efficient and reliable silencing of cortical networks, we recorded calcium activity of layer 2/3 pyramidal cells while stimulating ChR2-expressing PV interneurons with blue light (Supplementary Figure 2a,b). Two-photon imaging was performed as indicated in ‘Longitudinal two-photon calcium imaging during learning’, except that only one plane was recorded (15.49Hz, 150-250*µ*m below dura, 312×192*µ*m^2^, x1.7 or x2 magnification, laser power *≤*40 mW). A mounted LED (490nm, M490L4, Thorlabs) and a LED driver (Thorlab, LEDD1B) were used to deliver blue light at six different power levels over the AC. Pure tones (4-64kHz, 80dB SPL) and complex sounds were played (100-ms duration each, 100-frame intervals) and blue light was delivered in a counterbalanced manner. On a silencing trial, a trigger command is sent 100ms before sound onset from Clampex to the Tower electronics (Scanbox) that generates control signals for the LED and the PMT shutter (LED on for 1ms, PMTs off for 9ms, repeat for 5 frames; Supplementary Figure 2c). The first pulse was triggered 68ms before the onset of the sound, and the stimulation continued for a total of 320ms (Supplementary Figure 2c). To estimate the LED powers at the cortical surface (in mW/mm^2^), we measured the LED power coming out of the objective and estimated the cortical surface illuminated to be 2 mm (16X Nikon objective), leading to LED powers ranging from 0 to 3.15 mW/mm^2^.

Non-rigid registration and cell segmentation were performed using suite2p^79^ (https://github.com/MouseLand/suite2p). Fluorescence of each putative neuron (*n* = 454) was extracted and converted into *Δ*F/F by taking the mean activity as the baseline. We aligned neural responses to tone presentation, and quantified the effect of optogenetic silencing by comparing the mean activity of each neuron across all repetitions of sound presentations at different light powers (Supplementary Figure 2d,e). Only *Δ*F/F in frames immediately following light presentation were considered for quantification to avoid light contamination of the signal.

### Calcium imaging preprocessing

Upon acquisition, images were cropped (to remove artifact bands on plane 1 due to the electronically tunable lens) and converted to HDF5 files. Non-rigid registration (suite2p,^79^ https://github.com/MouseLand/suite2p) was run on the concatenated movie of all files recorded for a given mouse. All motion-corrected movies were visually inspected. Because recordings were made over weeks for a given dataset, our dataset could contain cells only weakly active overall. We, therefore, opted for manual detection of regions of interest (ROIs) rather than a semi-automatic one that uses cell activity to detect ROIs (e.g. suite2p cell registration). Manual ROI drawing was done in ImageJ using mean enhanced and maximum projection images. We identified 7,137 ROIs in 8 mice, with an average of 892*±*109 ROIs per mouse. The stability of each ROI throughout the entire recording was then assessed using a custom-written GUI in Matlab (MathWorks, Natick, MA). Overall, 2,332/3,935 cells were tracked every day of the task training in Learning mice (mean proportion of 67.3*±*7.5% of total ROIs per mouse), and 2,321/3,202 cells were tracked every day of passive exposure in Passive mice (mean proportion of 87.6*±*6.2% of total ROIs per mouse). Fluorescence activity from the ROIs was extracted using custom functions (Matlab). Raw fluorescence of each cells was then normalized as:

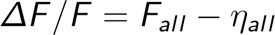

where

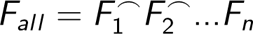

where the symbol ⌢ represents a concatenation, *n* is the number of files, 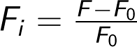 with *F* the raw fluorescence extracted from recording file *i* and *F*_0_ the median of this time series. *η_all_* is the median of *F_all_* over a sliding window of *∼*3 minutes. To compare calcium activity across trials, baseline fluorescence (activity during the inter-trial interval, before tone onset) was subtracted from the trial activity, so the *ΔF/F* reflected changes of intensity to the original intensity before trial onset.

### Data analysis

#### Statistics

Analyses were performed in Matlab (MathWorks, Natick, MA), using custom written programs, FMAToolbox (M. Zugaro, http://fmatoolbox.sourceforge.net), and Tensor Tool-box for MATLAB (https://www.tensortoolbox.org/). Descriptive statistics are reported as mean *±* standard error of the mean when the underlying distribution is Gaussian-shaped (Jarque-Bera test) or median *±* standard error of the median otherwise. Unless indicated otherwise, bars represent median *±* standard error of the median, box-plots represent median (center line), upper and lower quartiles (box limits) and 1.5x interquartile range (whiskers), and all statistical tests were two-sided. Student’s t-test was used for two group comparisons of Gaussian distributions, paired t-test for paired Gaussian distributions. For non-Gaussian distributions of independent data, two group comparisons were made using Wilcoxon rank sum tests. Wilcoxon sign rank tests were used for two group comparisons of non-Gaussian paired data or to compare medians of non-Gaussian distributions to single values. Two-way ANOVAs were performed to evaluate the effects of two independent variables on data and their interaction. All ANOVA statistics are reported in Supplementary Table 1. Proportions were compared using the binomial proportion test. Distributions were compared using the Kolmogorov-Smirnov test. No statistical methods were used to pre-determine sample sizes, but our sample sizes are similar to those generally employed in the field. Data collection and analysis were not performed blind to the conditions of the experiments.

#### Behavior analysis

Rare non-learner mice were excluded and massive drops in performance after reaching high performance (accuracy *>* 0.7) were not analyzed. Accuracy in probe and reinforced context was computed as (*n_HIT_* + *n_CR_*)*/*(*n_S_*_+_ + *n_S−_*), where *n_HIT_*, *n_CR_*, *n_S_*_+_, and *n_S−_* are the number of hit, correct reject, S+ and S*−* trials, respectively. To have trial-resolution assessment of behavior, we also computed response index curves (Fig. 1i), which reflected the latency to respond to the cues compared to local, spontaneous licking rate^6,80^. Response index curves were computed for the two cues (S+ and S*−* trials) separately as the latency to lick in a 2.5s window before the cue onset minus the latency to lick in the response window (2.5s after cue onset). If no lick was detected in either of these windows, the latency was set to the window duration, i.e. 2.5s. Therefore, for a given trial, the response index ranges from *−*2.5 to +2.5, with positive values indicating that the response to the cue was shorter than the local spontaneous licking rate of the animal, negative values indicating a decrease of licking in response to the cue, and values around 0 indicated that the cue did not impact the response rate. Performance index (Fig. 4s,t) was computed as the difference between S+ and S*−* cumulative response index curves. From the S+ response index, we identified the ‘change point’ (CP)^6,80^, i.e. the trial after which there is a consistent expression of cued behavior (Fig. 1i). We used the method described here^80^, itself a variation of the method used in^6^. Briefly, a recursive algorithm successively run over each data point *i* of the cumulative S+ response index curve and performs the following steps: 1) draws a straight line from trial *i* to trial 0 or the previous true CP, whatever is the closest to *i* and identifies the point that deviates maximally from this line as a putative CP; 2) calculates the strength of the evidence that it is a true CP, i.e. the log of the odds against the null hypothesis of no change (the logit). If logit *>* 1.3^6,80^, the putative CP becomes a true CP. As multiple CPs can be identified on a single curve, we reported in Fig. 1i only the first CP associated with a positive change of the slope of the cumulative behavioral responses^80^.

#### Best frequency

Single cell responses to the 17 tones presented were evaluated with paired t-test comparing pre- vs post-tone mean activity (over 10 frames, *∼*626ms). Bonferroni correction for the number of sounds (*n* = 17) was applied. For each cell, the peak amplitude response to each tone was determined as the maximum value of the averaged traces in the 10-frame post-tone window. A neuron’s best frequency was determined as the pure tone for which the peak amplitude response was the highest among significant responses only.

#### Tone-evoked responses across days

Evolution of tone-evoked responses in the reinforced context was analyzed using all cells recorded (Supplementary Figure 12 and Supplementary Figure 13) but the conclusions held when restricted to cells tracked every day. Response to S+ and S*−*, or stimulus 1 (S1) and stimulus 2 (S2) for Passive mice, were analyzed separately with paired t-tests comparing pre- vs post-tone mean activity (in 11-frame windows, *∼*688ms). A cell was considered tone-responsive in a given day if it significantly responded to either S+/S1 *or* S*−*/S2. Given that response profiles were identical to S1 and S2, responses to the two tones were sometimes represented together (Supplementary Figure 13).

#### Tone-evoked responses, responsiveness, response index and stimulus selectivity index

Tone-evoked responses were defined as the mean *Δ* F/F in a 11-frame window (*∼*688ms) post tone onset. Responsiveness was defined as the proportion of cells exhibiting a significant tone response (paired t-tests; Supplementary Figure 13). To compute response indices (Fig. 3k), the peak of the average *Δ* F/F for hit and S*−* trials (FA trials until mid-expression, CR trials after that) in 80-trial blocks was calculated, followed by the proportion of blocks with significant (peak *Δ* F/F *>* 2% of baseline) response throughout learning. The response index of a neuron was computed as the average response probability in hit and S*−* trials over learning. Stimulus selectivity was computed for each neuron in 80-trial blocks over learning and defined as:

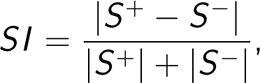

where *S* ^+^ is the peak *Δ* F/F in the tone-evoked response window on hit trials, *S^−^* is the peak *Δ* F/F in the tone-evoked response window on S*−* trials. SI could therefore ranged from 0 to 1, with 1 indicating maximal selectivity for either the S+ or the S*−*. Values of *S* ^+^ and *S^−^<* 2% were set to zero, and SI in blocks where *S* ^+^ and *S^−^* were both equal to zero was set to zero. The selectivity index of a neuron was its average SI over learning (Fig. 3l).

#### Stimulus decoding

For each mouse, cue identity was decoded across trial frames from activity of cells tracked across all days using linear discriminant analysis with 5-fold cross-validation (Supplementary Figure 1). Tone decoding accuracy in the tone-evoked window referred to the mean accuracy in the tone-evoked window (11 frames post-tone onset; Supplementary Figure 1e). Chance accuracy level was estimated by decoding cue identity across trial after randomly shuffling cue identity across trials (*n* = 20 shuffles/day/mouse).

#### Data organization and tensor decomposition

To analyze our high dimensional dataset, we took advantage of tensor decomposition^81–83^, a method that enables unbiased and interpretable descriptions of dynamic changes at multiple timescales, also referred as ‘tensor component analysis’ or TCA^26^. Here we used it not only to reveal within and across trial dynamics^26^, but also to identify shared and distinct variability in cell networks recorded from Learning and Passive mice. We organized calcium traces into a fourth-order tensor (or four-dimensional array) with four axes corresponding to individual neurons (recorded in Learning and Passive mice), time within trial, trials over time, and trial types. We then fit a tensor CANDECOMP/PARAFAC (CP) decomposition model^83–85^ to identify in an unsupervised way a set of low-dimensional components describing variability along each of these four axes (also referred here as factors; Supplementary Figure 9).

##### Data organization

We first built two arrays for learning and passive data separately and combined them afterwards. Only data from the reinforced context was taken for Learning mice. We filtered out disengagement periods (hit rate *≤* 0.5 in a 20-trial block), sometimes occurring during the last dozens of trials of the day and associated with significant changes in neuronal dynamics compared to engaged state^23,45,56–63^. For both Learning and Passive data, *Δ*F/F of each trial was selected from *−*1s to +4s relative to tone onset (2nd tensor dimension). With 4,643 cells tracked all days, 75 frames/trials, *∼*300 trials/day over 15 days, our dataset approximated 1,567,000,000 data points. To reduce computation time, trials of identical types (hit, miss, FA or CR) within 20-trial blocks were averaged together. In other words, from a given 20-trial block, up to four trial traces could be obtained (4th tensor dimension). Because of the exclusion of disengaged periods and the tendency of the animals to lick, miss trials were too rare in the Learning group to be considered without adding significant noise and were excluded. As a result, the 4th tensor dimension dissociated S+ (hit trials for Learning data, miss trials for Passive data), FA and CR trials. Finally, a crucial goal of this analysis was to be able to identify neural dynamics associated with task learning, and more precisely to isolate any dynamics associated with task contingency acquisition (measured in the probe context) or performance improvement (measured in the reinforced context). To this end, we aligned the trial traces to learning phases (3rd tensor dimension). First, we identified Acquisition, Expression and Expert phases in our 5 learning mice (see Supplementary Figure 8). The Acquisition phase started at the first trial of training and continued until maximum accuracy was reached in probe or when accuracy was *≥* 0.65 in probe and *≤* 0.70 in reinforced trials. This marked the beginning of Expression phase, which continued until Expert phase started at the second day of high and stable performance. Data in between Acquisition and Expert phases was part of the Expression phase. Evolution of individual mouse performance per identified phases is quantified in Supplementary Figure 8f. Resultant mega-mouse performance (i.e. pooled performance in 20-trial block across mice) is shown in Supplementary Figure 8d,e. Second, because these phases varied in duration across animals, we identified the mouse with the minimum number of trial traces in a given phases and downsampled the number of trial traces of the other mice to match this number. Down-sampling was performed by preserving the duration/performance range in each mouse (i.e. keeping first and last trial traces) and removing trial traces at consistent intervals in-between, such as the overall learning evolution of the phase was preserved. Third, each Passive mouse was assigned with the learning phases of a Learning mouse, and the same downsampling procedure was used. Finally, the two four-dimensional arrays containing Learning and Passive data, respectively, were concatenated in the first (neurons) dimension (referred as the ‘mega-mouse’ tensor) and *Δ* F/F traces were z-scored. Because Passive mice essentially did not lick, any data for FA trials for Passive cells were zeroed out. Any missing entries of the mega-mouse tensor were also zeroed out.

##### Tensor decomposition

To deal with incomplete data (absence of FA trials in Passive mice and possible missing CR early in learning or missing FA at expert level for Learning mice), we fitted an R-component weighted CP model^27^ to our mega-mouse tensor. Briefly, CP decomposition decomposes a tensor into a sum of rank-one tensors. For a third-order tensor *X ∈* R*^I×J×K^*, we wish to write it as:

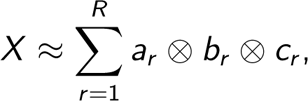

where *⊗* represents the vector outer product, *a_r_ ∈* ℝ*^I^*, *a_r_ ∈* ℝ*^I^* and *a_r_ ∈* ℝ*^I^* for *R* = 1, *. . . R*, and *a_r_ ⊗ b_r_ ⊗ c_r_* is a rank-one tensor. With perfect data we would obtain equality; however, in practice the presence of noise prevents it. We can use the Kruskal operator to simplify the previous expression^86,87^:

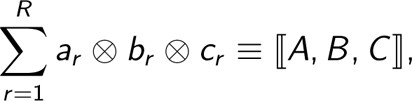

where factor matrices *A ∈* ℝ*^I×R^*, *B ∈* ℝ*^J×R^* and *C ∈* ℝ*^K×R^*, with

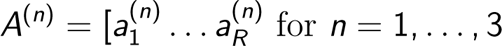

To fit the CP decomposition model to data, we used the CP-WOPT (CP Weighted OPTimization) algorithm^27^ that uses a first-order optimization approach to solve the weighted least squares problem, i.e. minimize the error function

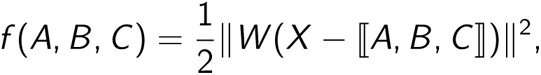

where *W* is a nonnegative weight tensor with same size as *X* defined as

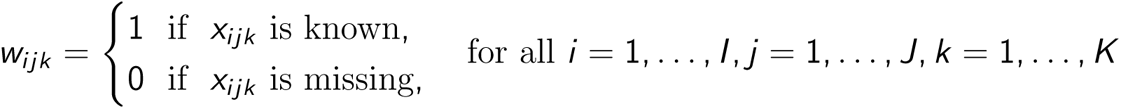

The weighted least squares objective function is solved over all the factor matrices simultaneously.

In practice, the rank *R* of a tensor is generally not known and is not easily determined^88^. To fit the CP models and choose the number of components, we closely followed the pipeline detailed in^26^. Briefly, we ran models 20 times with different random initializations for different numbers of low-dimensional components *R* = 1, *. . .*, 6. We used two metrics to compare and assess models: 1) the (normalized) weighted squared reconstruction error, computed for each fitted model, defined as:

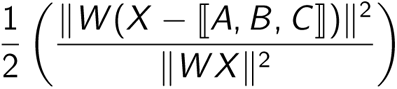

and 2) a similarity score^26,89^, quantifying the match between two fitted models i.e. how similar are the components resulting from two different runs. Let’s consider the Kruskal form of the tensor *X* (or ktensor)

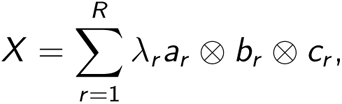

where *λ_r_* is the scaling factor after rescaling *a_r_*, *b_r_* and *c_r_*to be unit length. Considering two tensors [*A*, *B*, *C*] and [*D*, *E*, *F*],

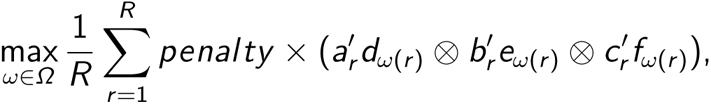

With

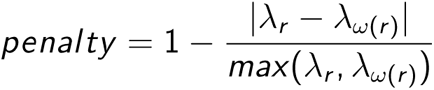

where *Ω* is the set of all permutations of the R components, and *ω* a particular permutation. With increasing number of components *R*, considering all possible matches is exponentially expensive and can be computationally prohibitive and factors were matched in a greedy fashion to identify good alignment (although not necessarily optimal). Similarity for each model fit was computed with respect to the best-fit model with the same number of components. Adding more components caused models to be less reliably identified (lower similarity score). For a given number of components *R*, the model fits were also visually inspected and compared. With our dataset, models with similarity scores above 0.8 were qualitatively similar while consistency dropped for values closed to 0.5. Therefore, a decomposition into 4 components was chosen for our dataset. The output of our decomposition was therefore a set of four components, each composed of four factors (i.e. weight vectors): 1) neuron factor (*W_N_*), reflecting cell ensembles, 2) within trial factor (*W_W_*), indicating when the activity occur in the trial, 3) across trial factor (*W_A_*), reflecting the evolution profile over learning/time at trial resolution, and 4) outcomes factor (*W_O_*), reflecting contribution of sensory, motor and cognitive variables. When *R* is small, increasing number of components demixed the activity until providing redundant information (when *R >* 4 for this tensor). Importantly, other types of decomposition were run, and other tensors (individual mouse, Passive and Learning data separately) were decomposed, and they all converged into the same description of the data.

#### Identification of learning-related dynamics

##### Quantification

To determine whether the low-dimensional dynamics described by the tensor decomposition were selectively attributed to the cells from Learning or Passive mice, we analyzed the neuronal factor, i.e. the neuronal weigths (*W_N_*) of the four components. We first compared the contribution of Learning and Passive networks to the highest (absolute) neuronal weights across components (Fig. 10c, Supplementary Figure 11c). Given that no constraint was applied on the sign of the weights, a given component could describe up to two distinct dynamics. We therefore also analyzed positive and negative neuronal weights separately (Supplementary Figure 10d,e, Supplementary Figure 11d) and obtained the same results: components 1 and 2 described dynamics largely driven by the passive network while components 3 and 4 described neural dynamics driven by the learning network. Importantly, we verified that this effect was not driven only by one mouse: for each component, we compared the neuronal weights of cell populations recorded in each mouse of a group (e.g. passive) and compared it to the other group (e.g. learning) (Supplementary Figure 10e). Because the components described different neuronal dynamics, this result therefore implied that learning and passive networks contained different low-dimensional dynamics.

##### Visualization

To visualize how the revealed neural dynamics maps onto our two experimental groups (learning and passive), we used two different dimensionality reduction approaches to project the data into a two- or three-dimensional space. First, we used t-distributed stochastic neighbor embedding (t-SNE) on the neuronal weight matrix *W_N_* of size *N × R*, where *N* is the number of cells in tensor and *R* the number of components (Fig. 2l,m, Supplementary Figure 10f). Second, we used principal component analysis (PCA) on different combinations of factors: *W_N_ ⊗W_W_* (Supplementary Figure 10g), *W_N_ ⊗W_W_ ⊗W_A_* (Fig. 2k), *W_N_ ⊗W_W_ ⊗W_O_* (Supplementary Figure 10h), and *W_N_ ⊗ W_W_ ⊗ W_A_ ⊗ W_O_* (Supplementary Figure 10i), and projected learning and passive data separately into the same principal component subspace.

#### Unique participation: defining cell ensembles

For visualization and quantification purposes, we attributed each neural dynamic to unique cell ensembles based on neurons’ weights (Supplementary Figure 11a). As indicated earlier, factor weights could be positive or negative and therefore up to two distinct dynamics could be represented per component. With this in mind, each neuron *i* was associated with a two digit code [componentID sign], i.e. a unique dynamic, where componentID is the component where the *|W_N_ |* of the neuron *i* was maximal. This approach therefore filtered out non-participating (i.e. low weighted) neurons in describing neuronal dynamics, as illustrated in Supplementary Figure 11b. Finally, in order to assess the nature of encoding of these cell ensembles, cell ensembles 1 and 2 were restricted to cells recorded in the passive mice, while cell ensembles 3 to 6, describing dynamics of components 3 and 4, were restricted to cells recorded in learning mice (Fig. 2m).

#### Comparison of calcium responses between trial outcomes with a time-changing signal

For each *Δ*F/F comparison between different trial types, both the number of trials taken (‘how many’) and the trial numbers (‘when’) were matched between group to control for time/learning effect and power/noise difference (Figs. 4d-h,m-n, 5f,g).

#### Analysis of licks outside task events

Lick bouts outside task events were defined as lick bouts that preceded the first tone presentation at the beginning of each behavioral block. The analysis was restricted to the first day of training, to remove learning confound as much as possible (Supplementary Figure 15b,c). A lick bout was defined as a succession of at least 3 licks with less than 1s interval in-between each lick. In addition, it had to be preceded by a 1s no lick period, used to z-score the traces.

#### Classification of false alarm trials based on reward prediction activity

For each learning mouse, we trained a two-class support vector machine (SVM) algorithm to decode trial identity (matched hit and CR trials) from late-in-trial activity (single trial AUCs) of neurons part of cell ensemble 5. This decoding gave us access to a misclassification rate (for each class and global), representing the noise level in the data (Supplementary Figure 16a,b,e). We then used this trained SVM to classify FA trials, reasoning that if a reward prediction signal is present during an FA trial, it will be decoded as a hit trial. In each mouse, the proportion of FA trials with a RP signal was higher than the misclassification rate of the decoder (Supplementary Figure 16e).

#### Isolating brief disengagement periods during behavior

Once mice acquire task contingencies and start increasing their correct rejection in the reinforced context, they generally stop behaving in the probe context (hit rate close to zero; e.g. Supplementary Figure 8)^7^. We therefore found these periods by looking for probe blocks with hit rate *<* 0.4 (Fig. 5g).

#### Pre- vs post-behavior changes in tonopy

To assess how learning and passive exposure affected the cortical tonotopic map, we compared best frequency surfaces from tuning curve recording sessions before and after learning (see ‘Pre- and post-task tonotopic mapping’). We first split the field of views in 30 *×* 30 pixels (*∼*41 *×* 41*µ*m) and computed the best frequency mode of the local neuronal population in each of those pixel blocks (Fig. 3o). We estimated the change in surface before and after behavior as:

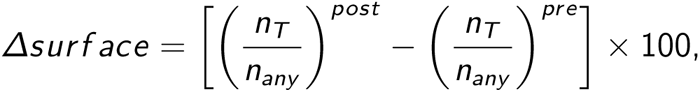

where *n_T_*is the number of pixel block with *T* best frequency mode and *n_any_*the number of pixel block with any best frequency. In our analysis, *T* could be the S+, S*−*, tones in between S+ and S*−*, and tones with lower or higher frequency than S+ or S*−* (Fig. 3n). We also evaluated best frequency mode differences before and after behavior in pixel blocks (Fig. 3q).

#### Spatial clustering of contingency-related cell ensembles

To assess the spatial distribution of reward prediction and action suppression cell ensembles (referred to here as ‘clusters’), we compared the distance between the two ensembles to a random spatial organization (Fig. 6a,b). To do so, we computed the median of between-cluster cell distances and compared it to a median distribution obtained with cell ensemble identity shuffles (*n* = 500). This allowed us to assess the clustered nature of these two cell ensembles while preserving the spatial cell distribution in the fields of view. We considered the cell ensembles significantly clustered if the median distance of the cell ensembles was *>* 97.5% of the shuffle distribution. Because of the different statistics of cell distribution inside a field of view for each mouse, comparing raw cell ensembles distances between mice was prohibited. Instead, we computed a z-scored distance for each mouse by subtracting the mean and dividing by the standard deviation of the shuffle distribution to the data median distance (Fig. 6c).

#### Pre-task stimulus selectivity index

For cells with positive tone-evoked responses to both S+ and S*−* in pre-task tuning curve session, pre-task stimulus index (SI, (Fig. 6d,e) was computed as:

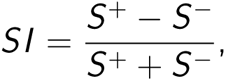

where *S* ^+^ is the peak *Δ*F/F in the tone-evoked response window to the S+ tone and *S^−^* is the peak *Δ*F/F in the tone-evoked response window to the S*−* tone. SI could therefore ranged from *−*1 to 1, with 1 indicating total selectivity for the S+, *−*1 indicating total selectivity for the S*−*, and zero an absence of selectivity (similar response to both tone).

#### Assessing the relationship between tonotopic map and contingency organization

To assess whether reward prediction cells were S+ preferring cells and action suppression cells were S*−* preferring cells before training started, we generated two separate statistical tests (Fig. 6g,h). First, we tested the hypothesis that the reward prediction cell ensemble emerged from S+ preferring cells. We constructed a distribution of best-frequency distance to S+ if H0 was true, i.e. if reward prediction cells were to have a best frequency the closest to S+ given the field of view statistics (Fig. 6g). Separately, we tested the hypothesis that the action suppression cell ensemble emerged from S*−* preferring cells. We constructed a distribution of best-frequency distance to S-if H0 was true, i.e. if action suppression cells were to have a best frequency the closest to S-given the field of view statistics (Fig. 6h). Finally, we compared the proportion of S+ and S-preferring cells among reward prediction and action suppression cell ensembles and observed no differences (Fig. 6i).

## Data availability

The data that support the findings of this study are available from the corresponding authors upon request.

---

**Supplementary Figure 1.**
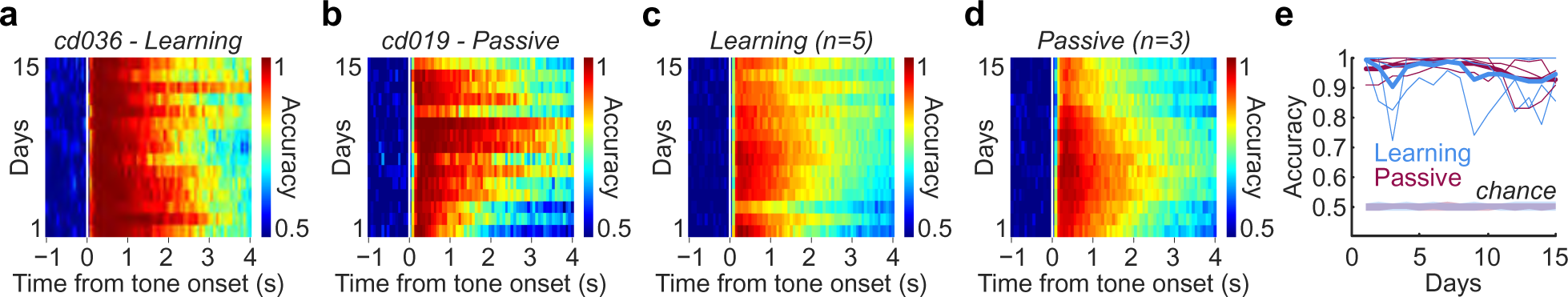
Stimulus decoding in the auditory cortex is at ceiling from Day 1 of learning. **a**, Stimulus decoding is at ceiling on Day 1 and remains high throughout learning (example mouse) Only the cells tracked across all days were used to decode tone identity. **b**, Stimulus decoding is at ceiling on day 1 and remains high throughout passive exposure over 15 days (example mouse). **c**, Average decoding accuracy for all Learning mice (*n* = 5). **d**, Average decoding accuracy for all Passive mice (*n* = 3). **e**, Evolution of tone decoding accuracy in the tone-evoked window across days for Learning and Passive mice compared to chance level (trial shuffle, see Methods).

**Supplementary Figure 2.**
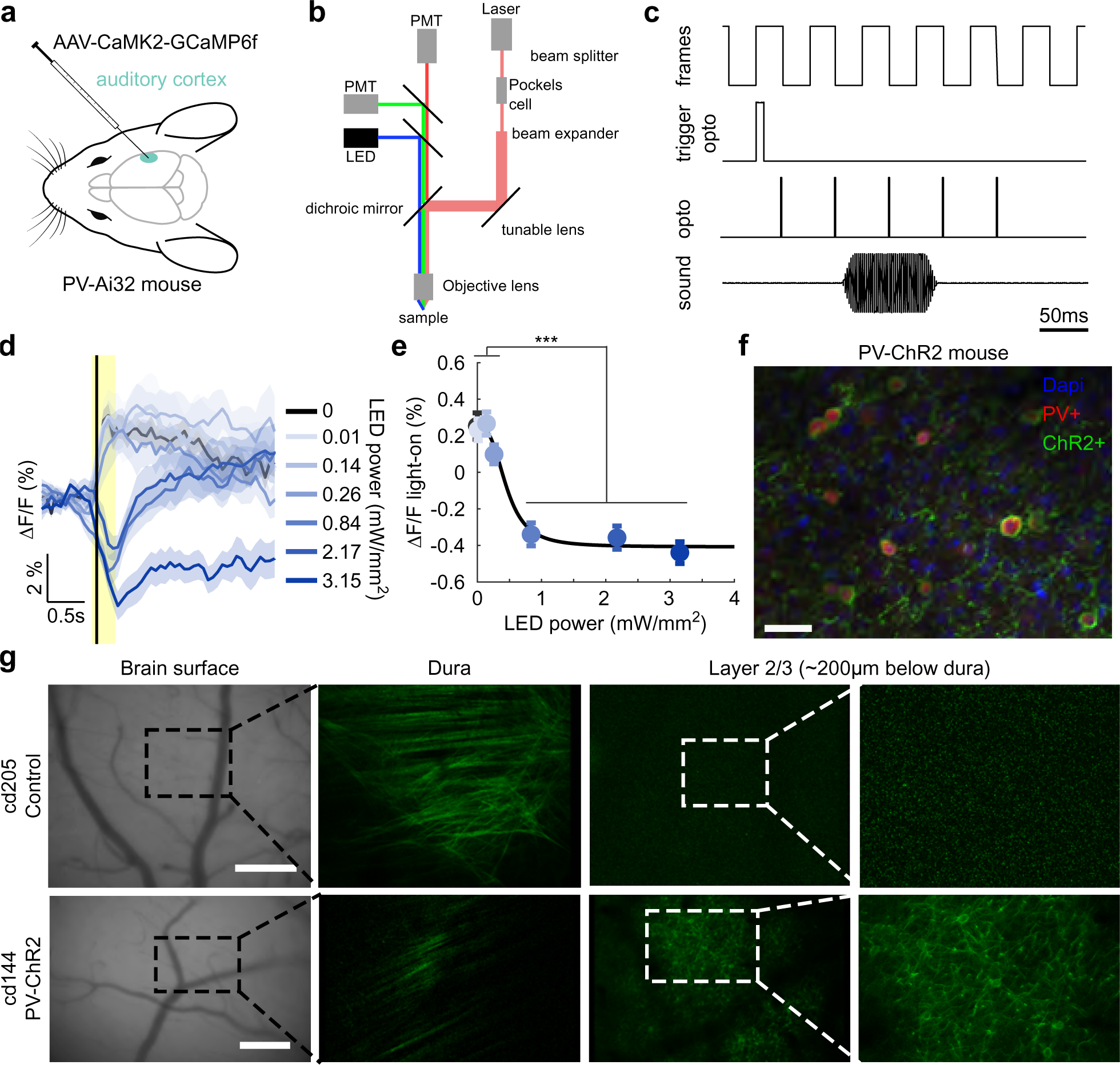
Activating PV+ neurons in the auditory cortex robustly suppresses stimulus-evoked activity of excitatory neurons. **a**, PV-ChR2 mice (*n* = 2) were injected with AAV-CaMKII-GCaMP6f to allow simultaneous one-photon excitation of PV cells and two-photon recordings of pyramidal cell population. **b**, Schematic of simultaneous widefield optogenetics and two-photon imaging. **c**, Optogenetic activation was locked to frame acquisition. **d**, Trial-averaged *Δ*F/F aligned to tone onset (black vertical line) of an example neuron at different intensity of LED power (blue scale). Yellow rectangle indicates period of light delivery. mean *±* s.e.m. **e**, Effect of optogenetic silencing as a function of LED power (*n* = 454 neurons; Friedman test, *p ∼* 0). *Δ*F/F at powers 0-0.26 mW/mm^2^ are all significantly different from *Δ*F/F at powers 0.84-3.15 mW/mm^2^ (post hoc comparisons with Tukey-Kramer test, \*\*\**p <* 0.001). Black line is the logistic fit. median *±* s.e.median. **f**, Immunostaining of PV-ChR2 mice auditory cortex showing ChR2 expression in PV cells (PV+ and ChR2+ colocalization). **g**, Post-task imaging of a representative control (top) and a representative test (PV-ChR2, bottom) mouse used in AC silencing experiments. Note that no fluorescence below the dura is detected in control mice.

**Supplementary Figure 3.**
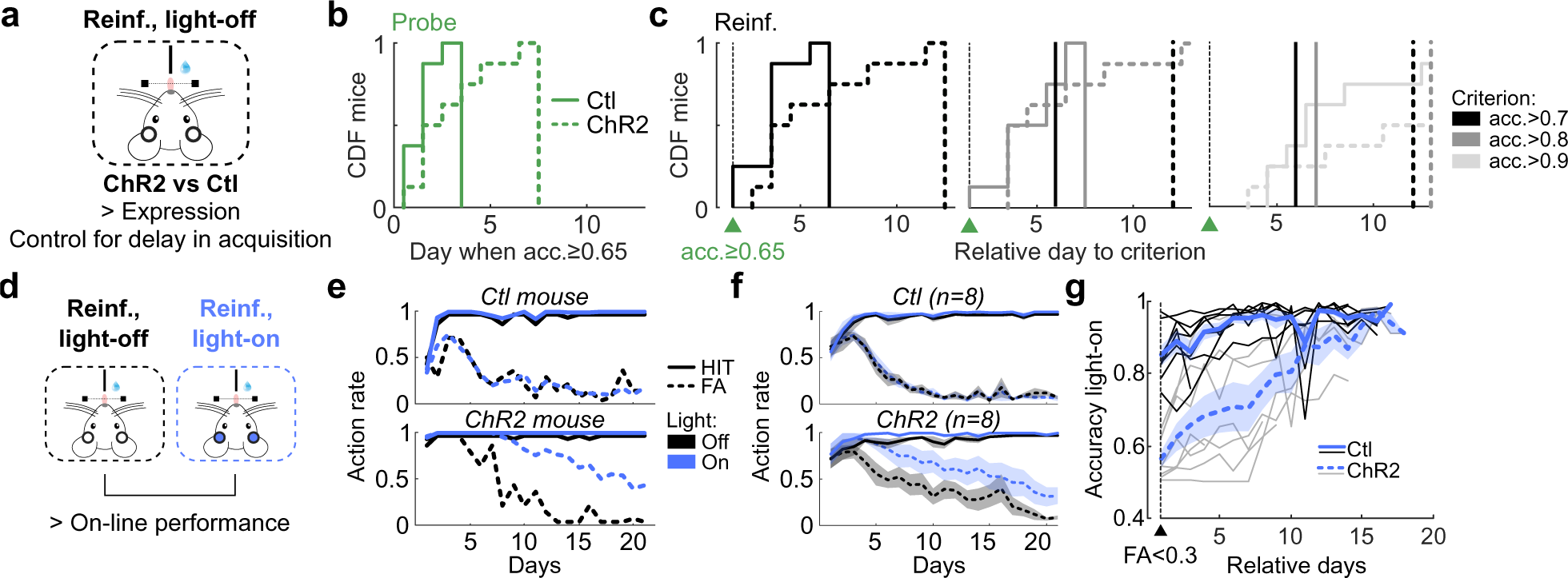
AC full trial silencing impairs expression and on-line performance. **a**, Assessment of the impact of AC full trial silencing over learning on Expression by controlling for the delay in Acquisition. **b**, Cumulative distribution function (CDF) of mice as function of the day to reach an accuracy *≥*0.65 in probe trials. **c**, Cumulative distribution function (CDF) of mice as function of the relative number of days to reach accuracy (acc.) criteria of *>*0.7 (left), *>*0.8 (middle), and *>*0.9 (right) in reinforced light-off trials after reaching an accuracy *≥*0.65 in probe trials. Black and dark gray vertical lines correspond to when CDF was reach for acc.*>*0.7 and *>*0.8, respectively. **d**, Comparing action rate and accuracy between reinforced light-off versus reinforced light-on trials to assess the impact of AC silencing on on-line performance. **e**, Hit (solid line) and FA (dashed line) of an example control mouse (top) and an example PV-ChR2 mouse (bottom) in reinforced light-off (black) and reinforced light-on (blue) trials across learning. **f**, Averaged action rate in reinforced light-off (black) and reinforced light-on (blue) trials per day for control (top) and PV-ChR2 (bottom) groups. **g**, Accuracy in light-on reinforced trials from the day when FA*<*0.3 in light-off reinforced trials. Note how PV-ChR2 mice (gray lines) increase accuracy (positive slopes) with light-on, showing that performance impairment fades away.

**Supplementary Figure 4.**
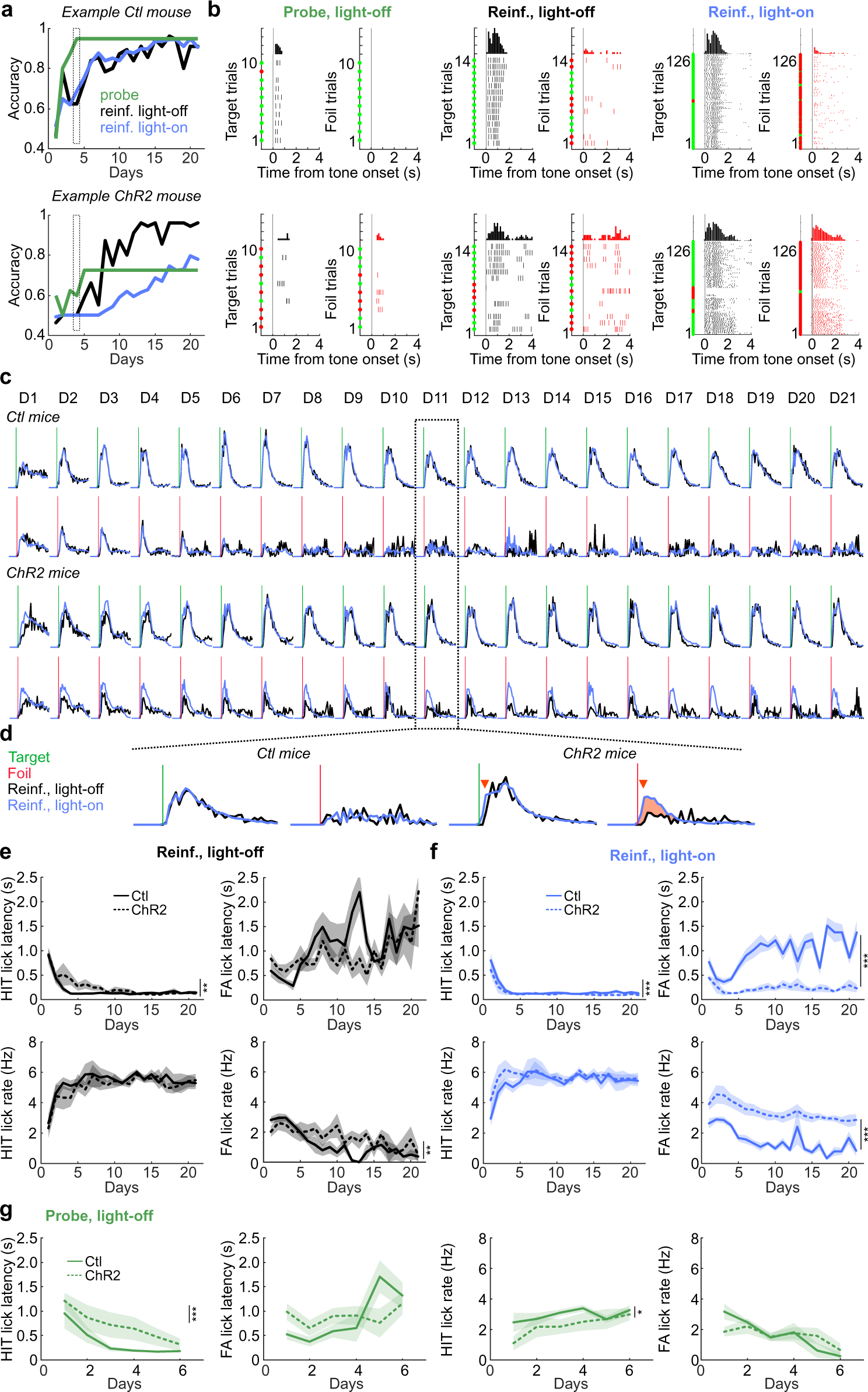
Effect of AC full trial silencing on lick patterns. **a**, Example control (top) and ChR2 (bottom) mice accuracy in probe light-off, reinforced light-off and reinforced light-on trials across day. Dashed rectangle indicates day where licks in **b** are extracted from. **b**, Lick raster plots from day 4 from the example mouse from **A** in probe light-off (left), reinforced light-off (middle) and reinforced light-on (right) trials, split into target (black, left) and foil (red, right) trials. Green and red dots indicates correct and incorrect trials, respectively. Note the difference in discrimination in all contexts between control and PV-ChR2 mice. **c**, Average lick probability across training days for control (*n* = 8) and ChR2 (*n* = 8) mice in response to target (vertical green line) and foil (vertical red line) tones, in reinforced light-off (black) and light-on (blue) trials. **d**, Insets showing faster lick latencies (red arrow heads) in response to both tones and higher lick probability in response to the foil (incorrect licking) in reinforced light-on compared to light-off in ChR2 mice (right). Light has no effect on lick structure in control mice (left). **e**, Lick latencies (top) and lick rate (bottom) in response to target (HIT trials; left) and foil (false alarm (FA) trials; right) tones in reinforced light-off trials (HIT lick latencies, Days: *F* (20, 256) = 8.2738, *p <* 10*^−^*^17^, Groups: *F* (1, 256) = 8.1568, *p* = 0.0046, Days*Groups: *F* (20, 256) = 0.9176, *p* = 0.56; FA Lick latencies, Days: *F* (20, 190) = 2.2393, *p* = 0.0027, Groups: *F* (1, 190) = 1.8422, *p* = 0.18, Days*Group: *F* (20, 190) = 1.5563, *p* = 0.067; HIT lick rate, Days: *F* (20, 256) = 4.3619, *p <* 10*^−^*^8^, Groups: *F* (1, 256) = 2.9549, *p* = 0.087, Days*Groups: *F* (20, 256) = 0.2927, *p* = 0.99; FA lick rate, Days: *F* (20, 190) = 4.04477, *p <* 10*^−^*^6^, Groups: *F* (1, 190) = 7.4070, *p* = 0.0071, Days*Groups: *F* (20, 190) = 1.1944, *p* = 0.26). **f**, Lick latencies (top) and lick rate (bottom) in response to target (HIT trials; left) and foil (false alarm (FA) trials; right) tones in reinforced light-on trials (HIT lick latencies, Days: *F* (20, 256) = 10.5303, *p <* 10*^−^*^22^, Groups: *F* (1, 256) = 11.2328, *p <* 10*^−^*^3^, Days*Groups: *F* (20, 256) = 0.6211, *p* = 0.90; FA Lick latencies, Days: *F* (20, 254) = 3.9111, *p <* 10*^−^*^6^, Groups: *F* (1, 254) = 450.4358, *p <* 10*^−^*^57^, Days*Group: *F* (20, 254) = 2.1947, *p* = 0.0029; HIT lick rate, Days: *F* (20, 256) = 2.6372, *p <* 10*^−^*^3^, Groups: *F* (1, 256) = 3.7748, *p* = 0.0531, Days*Groups: *F* (20, 256) = 0.4520, *p* = 0.98; FA lick rate, Days: *F* (20, 254) = 6.4469, *p <* 10*^−^*^13^, Groups: *F* (1, 254) = 301.2679, *p <* 10*^−^*^44^, Days*Groups: *F* (20, 254) = 0.6326, *p* = 0.89). **g**, Lick latencies (left) and lick rate (right) in response to target (HIT) and foil (FA) tones in probe light-off trials (HIT lick latencies, Days: *F* (5, 83) = 6.4522, *p <* 10*^−^*^4^, Groups: *F* (1, 83) = 11.7734, *p <* 10*^−^*^3^, Days*Groups: *F* (5, 83) = 0.2878, *p* = 0.92; FA Lick latencies, Days: *F* (5, 58) = 2.9217, *p* = 0.020, Groups: *F* (1, 58) = 0.9337, *p* = 0.338, Days*Group: *F* (5, 58) = 2.1909, *p* = 0.068; HIT lick rate, Days: *F* (5, 83) = 2.0103, *p* = 0.086, Groups: *F* (1, 83) = 5.9422, *p* = 0.017, Days*Groups: *F* (5, 83) = 0.5721, *p* = 0.72; FA lick rate, Days: *F* (5, 58) = 5.6386, *p <* 10*^−^*^3^, Groups: *F* (1, 58) = 0.0192, *p* = 0.89, Days*Groups: *F* (5, 58) = 1.6182, *p* = 0.17).

**Supplementary Figure 5.**
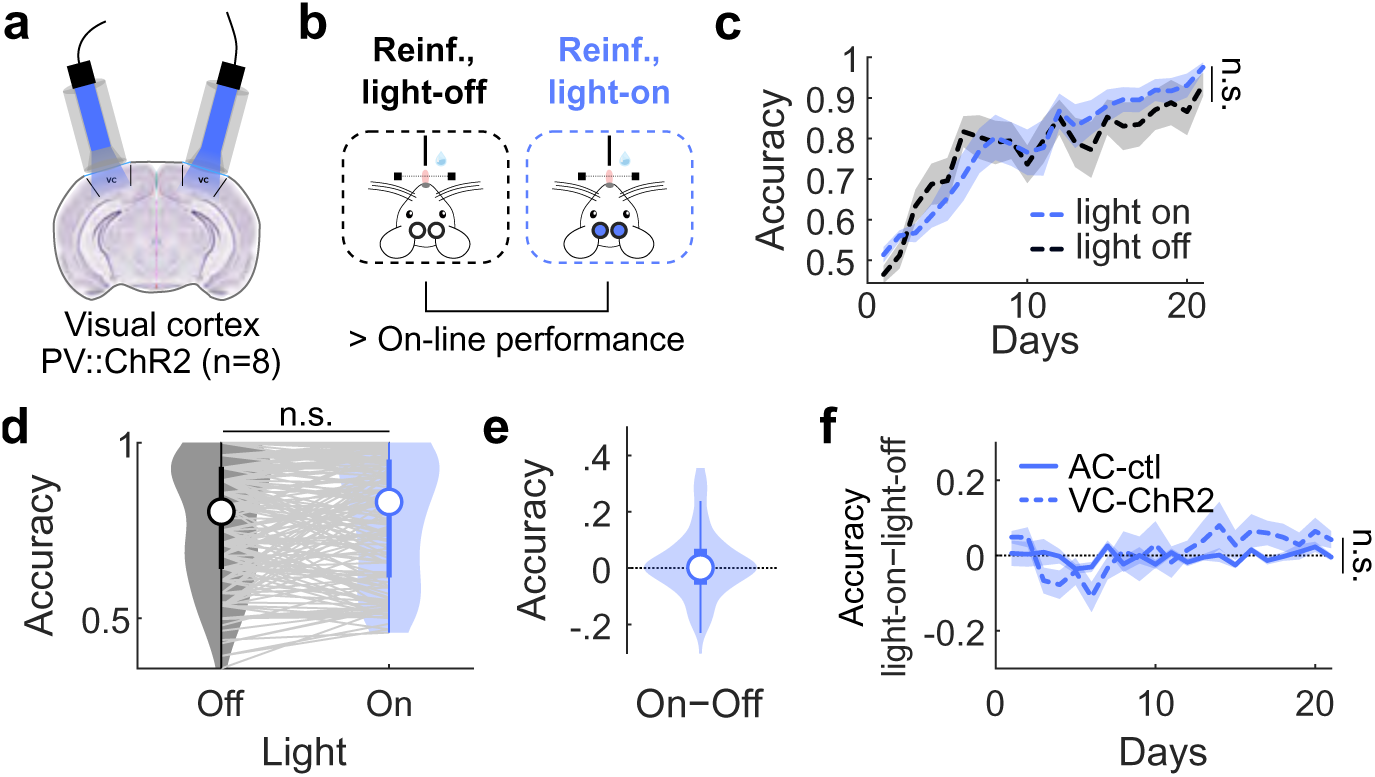
Silencing of the visual cortex does not impair performance throughout learning. **a**, Silencing of the visual cortex in 90% of the reinforced trials throughout learning (*n* = 8 PV-ChR2 mice). **b**, Comparison of reinforced light-off versus light-on trials shows no deficit when silencing the VC demonstrating the specificity of the effects of AC silencing. **c**, Accuracy in reinforced light-off and light-on trials across days (two-way repeated measures ANOVA, Group: *F* (1, 140) = 0.5093, *p* = 0.50). **d**, Accuracy in reinforced light-off and light-on trials (*n* = 168 sessions; Wilcoxon signed rank, *p* = 0.41). **e**, Difference in accuracy in reinforced light-on versus light-off trials per session. **f**, Difference in accuracy in reinforced light-on versus light-off trials across days in visual cortex PV-ChR2 mice (dashed line) versus auditory cortex control mice (solid line) (two-way ANOVA, Days: *F* (20, 271) = 1.5547, *p* = 0.06, Groups: *F* (1, 271) = 2.3072, *p* = 0.13, Days*Groups: *F* (20, 271) = 1.1540, *p* = 0.2950).

**Supplementary Figure 6.**
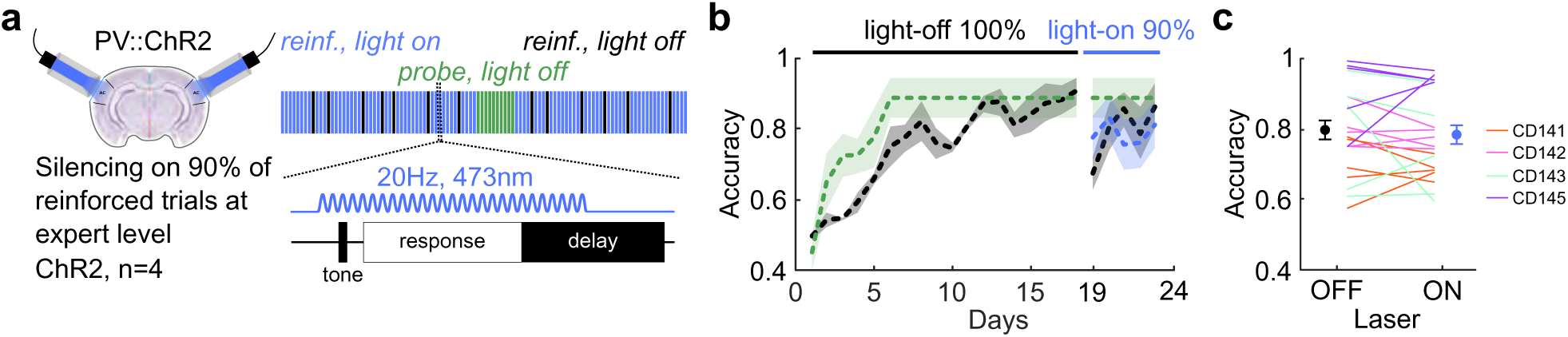
AC full trial silencing at expert level. **a**, Probabilistic optogenetic silencing of the auditory cortex at expert level. Silencing starts once stable performance is reached. **b**, Accuracy in probe light-off (green), reinforced light-off (black) and reinforced light-on (blue) trials. Silencing is performed from day 19 to 23. **c**, Accuracy in reinforced light-off and light-on trials (paired t-test, *p* = 0.602).

**Supplementary Figure 7.**
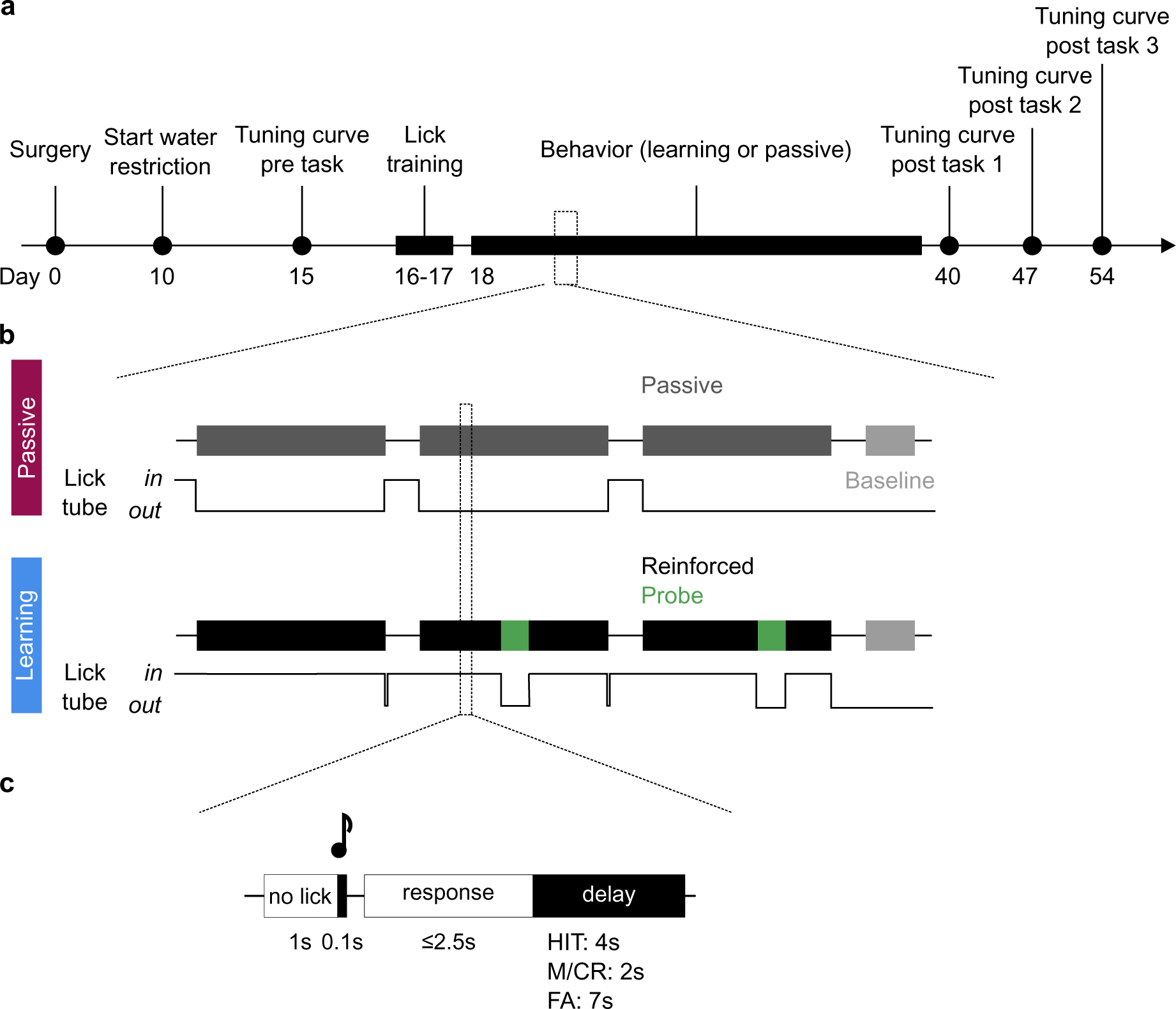
Experimental design and timeline of imaging experiments. **a**, After surgery, animals underwent a 10-day recovery period after which water restriction started. Tonotopic mapping (tuning curve session) of the auditory cortex took place 5 days later under the two-photon microscope, followed by two days of lick training under the two-photon microscope. These two sessions also allowed for habituation to head fixation and context. Behavior sessions started the following day for 15 or 16 days, after which tonotopic mapping sessions took place at day +1, +7 and +15 post learning. **b**, One behavioral session consisted of three blocks of 80 or 100 trials, and a baseline session (no tone presented). Two groups of mice were imaged under the two-photon microscope: the Passive group (top; *n* = 3) was presented with two pure tones but was never rewarded (lick tube out), and the Learning group (*n* = 5) was rewarded (3*µ*l water drop) if licking in the response window after the S+ tone. Two probe blocks of 10 trials each were introduced in two of the three reinforced blocks. **c**, Trial structure. After a no-lick period of 1s, a 100-ms tone was played, followed by a 200-ms dead period and a *≤*2.5s response period. The length of the delay period was of 2s after a miss (M, no lick after S+) or a correct reject (CR, no lick after S-), 4s after a hit (H, lick after S+) and 7s after a false alarm (FA, lick after S-).

**Supplementary Figure 8.**
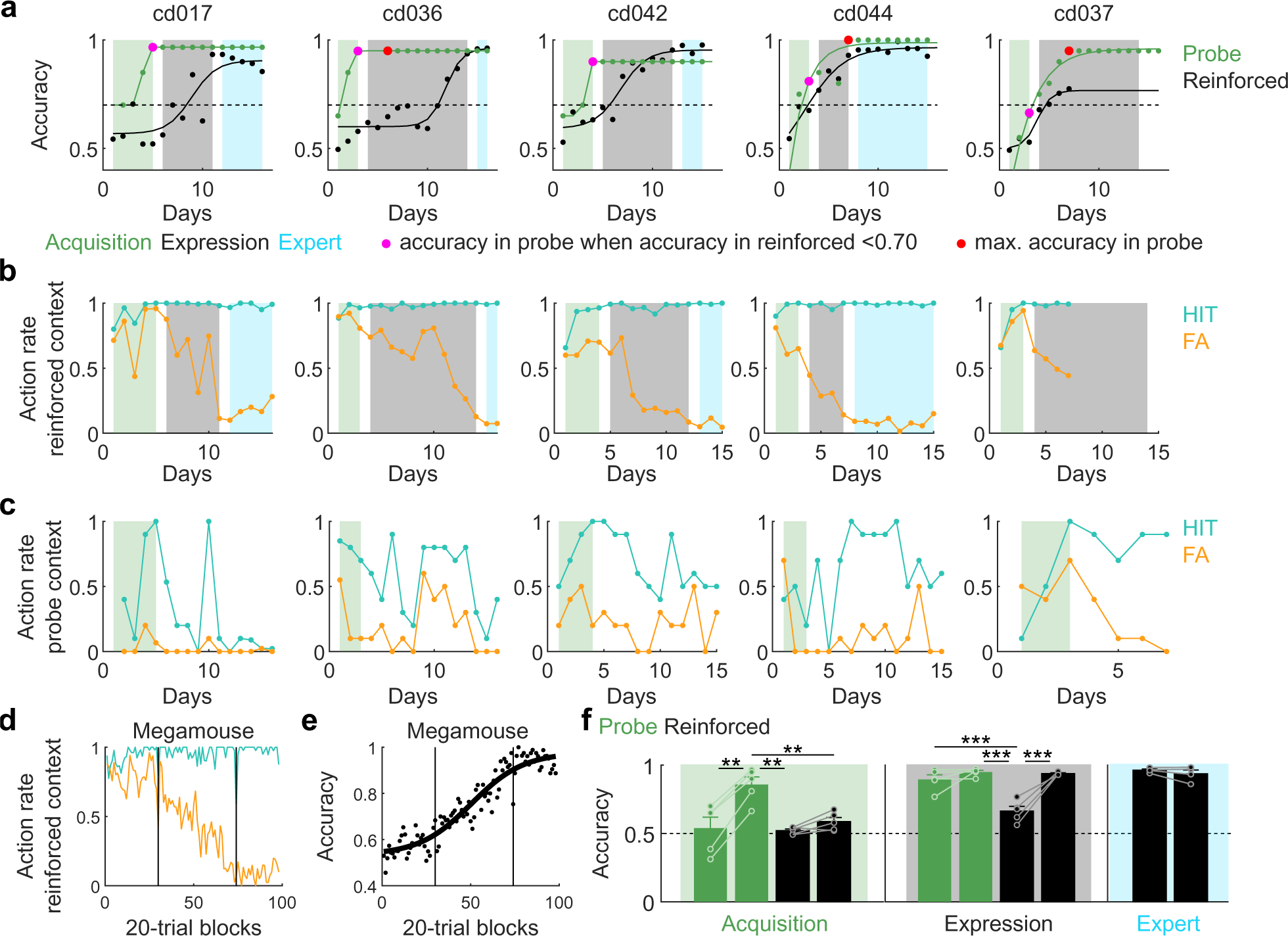
Inter-subject performance alignment for megamouse tensor. **a**, Accuracy in probe and reinforced contexts across days of all Learning mice. **b**, Action rate in reinforced context across days of all Learning mice. **c**, Action rate in probe context across days of all Learning mice. Please note that we fixed the probe performance at the maximum discrimination that was followed by a decrease in hit rate do to extinction. **d**, After the alignment procedure, action rate from the megamouse (all learning mice pooled) in reinforced context across learning phases. **e**, Megamouse accuracy in reinforced context across learning phases. **f**, Accuracy difference between the start and the end of the three learning phases in probe (green) and reinforced (black) contexts. Acquisition is characterized by an increase of accuracy in probe trials (paired t-test, *p* = 5.47.10*^−^*^4^) but not in reinforced trials (paired t-test, *p* = 0.07), Expression corresponds to an increase of accuracy in reinforced trials (paired t-test, *p* = 0.008) and Expert is when accuracy in reinforced trials is high and stable (paired t-test, *p* = 0.27).

**Supplementary Figure 9.**
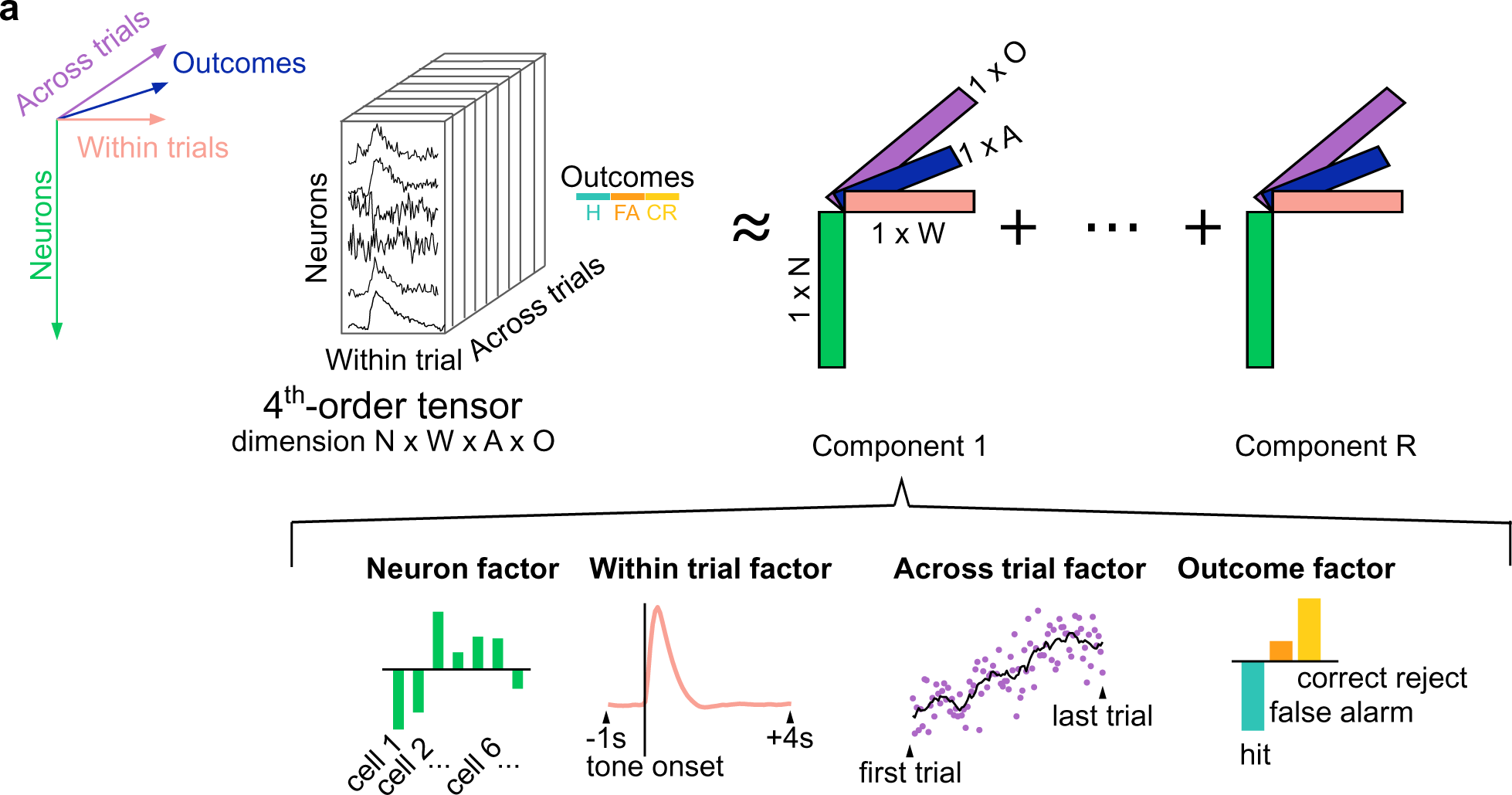
Tensor representation of neural data. **a**, Data are organized into a fourth-order tensor with dimensions N*×*W*×*A*×*O. Tensor decomposition approximates the data as a sum of outer products of four vectors. Each outer product contains a neuron factor (green rectangles), within trial factor (pink rectangles), across trial factor (blue rectangles) and outcome factor (purple rectangles). Each set of low-dimensional factors (i.e. component) describes the activity of group of neurons within and across trials according to trial outcomes.

**Supplementary Figure 10.**
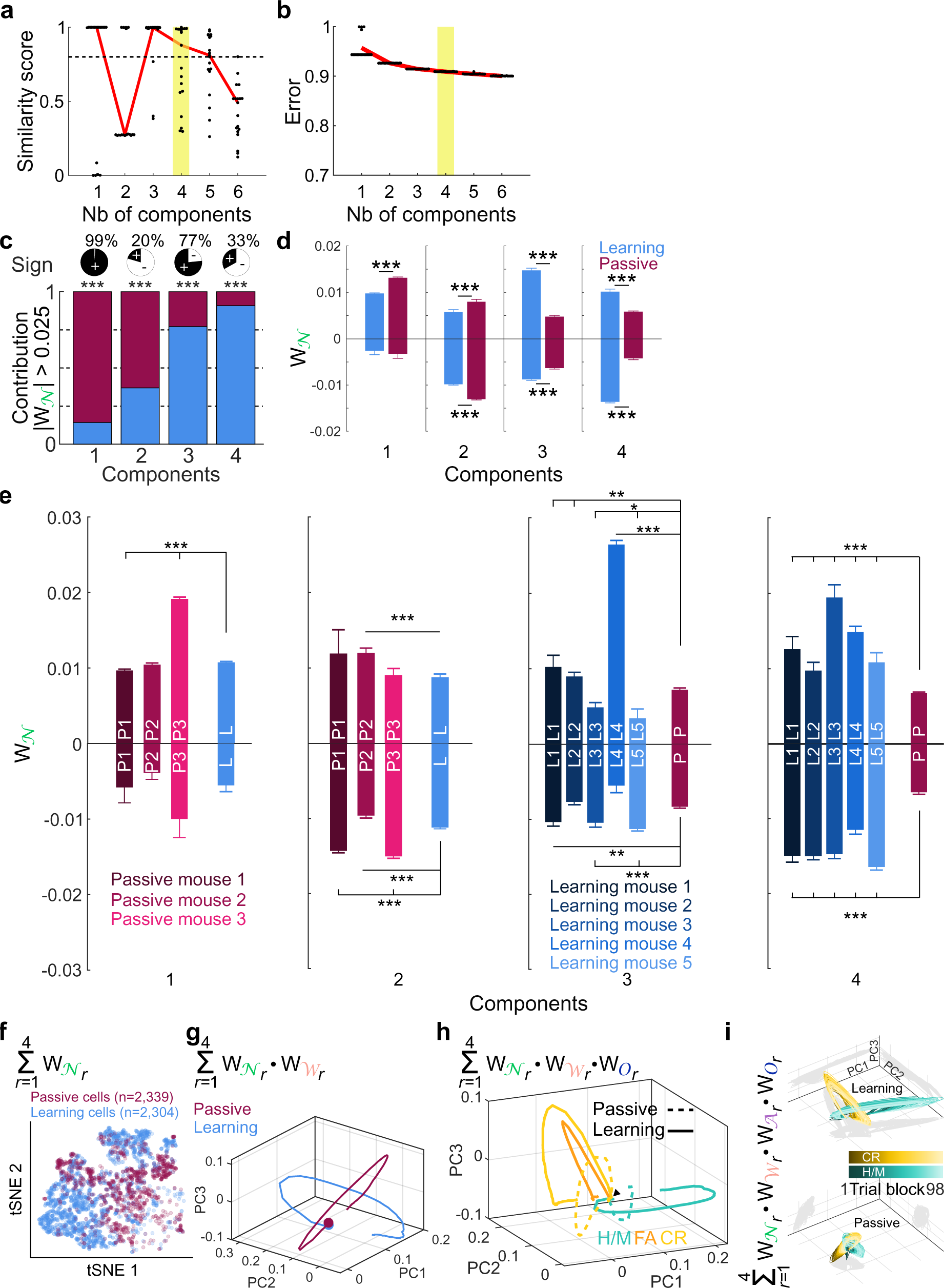
Low-rank tensor decomposition. **a**, Similarity score as a function of model components. Each dot shows the similarity of a single optimization run compared to the best-fit model within each category. **b**, Model reconstruction error as a function of the number of components, where each dot corresponds to a different optimization run. **c**, Neuronal contribution (Learning vs Passive cells) per components (binomial proportion tests, all *p <* 0.001). **d**, Positive and negative neuronal weights across components in cell population recorded in learning mice (Learning) or in passive mice (Passive) (Wilcoxon tests). **e**, Positive and negative neuronal weights across components and individual mice. **f**, t-SNE of neuronal weights. Note how Learning and Passive cell populations are largely non-overlapping. **g**, Projection of neuronal *×* within trial weights of Learning and Passive network activity into principal component space. **h**, Projection of neuronal *×* within trial *×* trial outcome weights of Learning and Passive network activity into principal component space. **i**, Projection of neuronal *×* within trial *×* across trials *×* trial outcome (H/M and CR only) weights of Learning and Passive network activity into principal component space.

**Supplementary Figure 11.**
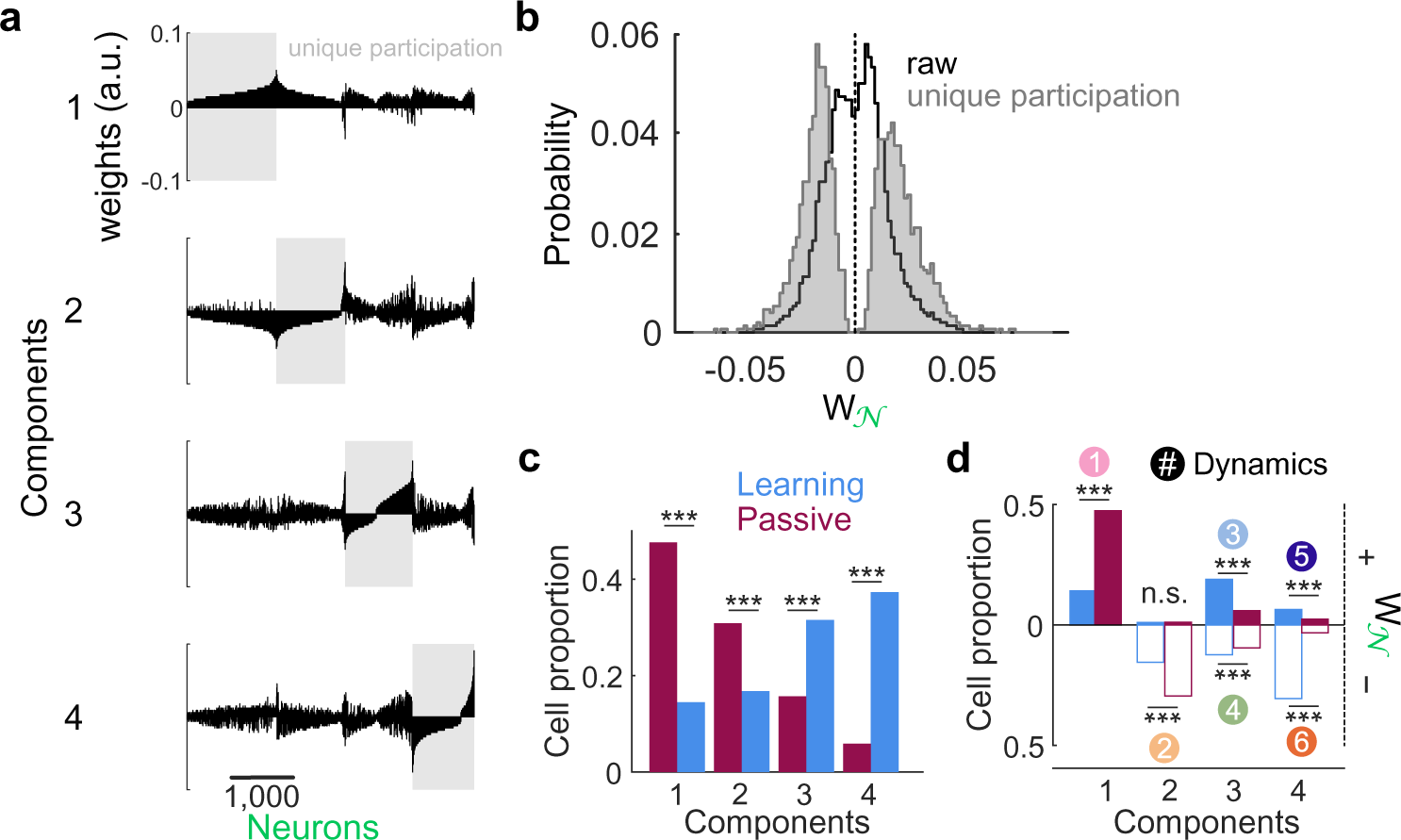
Defining unique cell ensembles based on neuronal weights. **a**, Neuronal weights in the four components. Each neuron is attributed to a given dynamic according to its highest absolute weights, i.e. highest contribution. As a result, each dynamic is attributed to a unique cell ensemble (gray rectangles). **b**, Neuronal weights distribution before (raw, black) and after unique contribution attribution (gray). **c**, Learning and Passive cell proportion among components after unique attribution (binomial proportion tests). **d**, Learning and Passive cell proportion among components and given neuronal weight sign after unique attribution. In other words, proportion of cells from Learning and Passive networks describing the tensor-revealed neuronal dynamics (binomial proportion tests). \*\*\**p <* 0.001, n.s.: not significant.

**Supplementary Figure 12.**
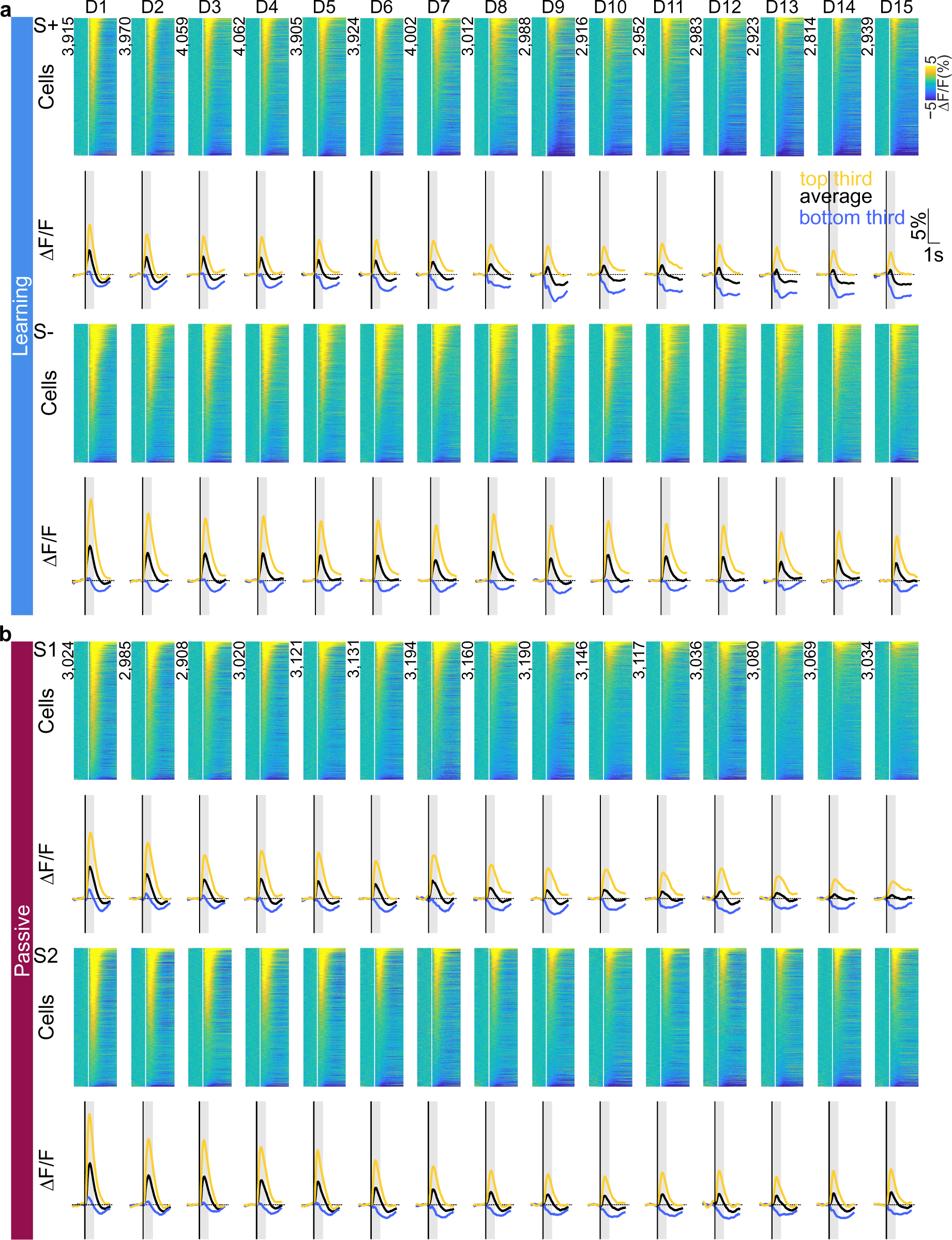
Evolution of tone-evoked responses across days. **a**, Tone-evoked responses to S+ and S*−* in Learning mice across days for all cells recorded. **b**, Tone-evoked responses to S1 and S2 in Passive mice across days for all cells recorded.

**Supplementary Figure 13.**
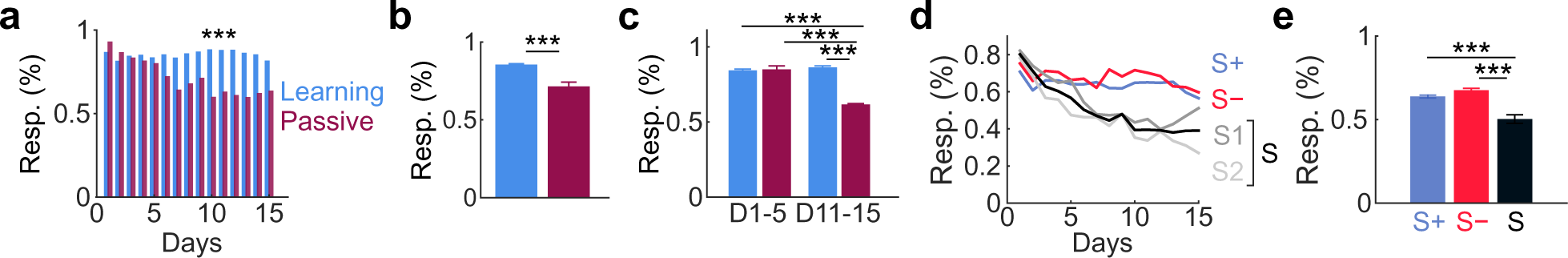
Learning counteracts tone-evoked habituation. **a**, Proportion of tone-responsive cells across days among Passive and Learning cells. **b**, Averaged proportion of tone-responsive cells in Passive and Learning networks (mean *±* s.e.m.; t-test, *p* = 3.89.10*^−^*^5^). **c**, Proportion of tone-responsive cells in days 1-5 versus days 11-15 in Learning and Passive networks (mean *±* s.e.m.; two-way ANOVA, Time *×* Group, *p* = 1.73.10*^−^*^7^). **d**, Proportion of cells responsive to S+ and S*−* in Learning network and S1, S2 or *S1 or S2* (S) in Passive network. **e**, Averaged proportion of cells responsive to S+, S*−* or S (mean *±* s.e.m.; ANOVA, *p* = 1.93.10*^−^*^6^).

**Supplementary Figure 14.**
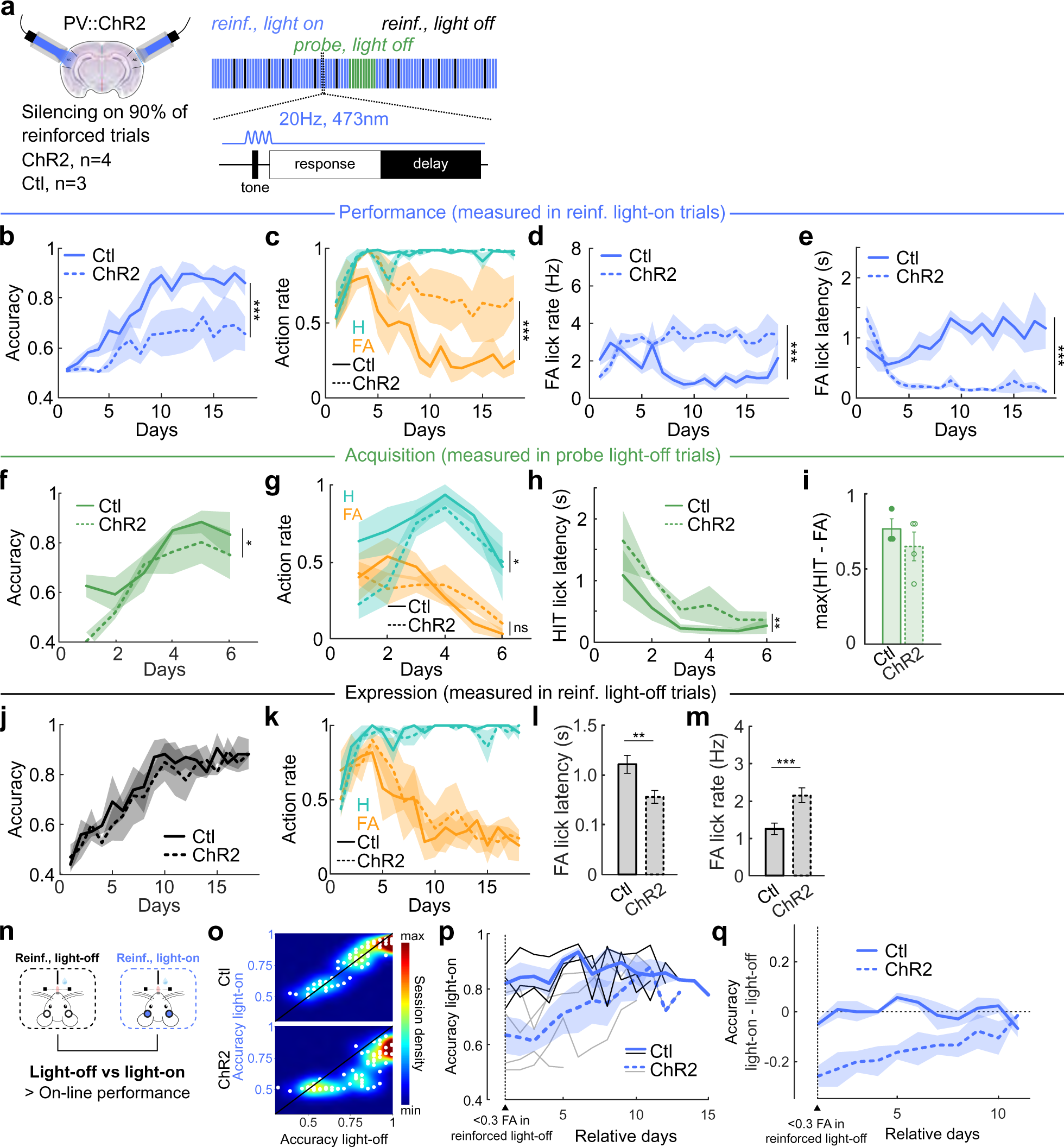
AC silencing restricted to sound presentation impairs audiomotor learning and on-line performance during learning. **a**, Probabilistic optogenetic silencing of the auditory cortex during learning. Light-on periods were restricted to sound presentation only (see Methods). **b**, Accuracy in reinforced light-on trials (two-way ANOVA, Days: *F* (17, 86) = 5.4950, *p <* 10*^−^*^7^; Groups: *F* (1, 86) = 50.5343, *p <* 10*^−^*^9^; Days*Groups: *F* (17, 86) = 0.70700, *p* = 0.79). **c**, Action rate in reinforced light-on trials (HIT, two-way ANOVAs, HIT, Days: *F* (17, 86)10.68010, *p <* 10*^−^*^14^; Groups: *F* (1, 86) = 0.0200, *p* = 0.89; Days*Groups: *F* (17, 86) = 1.0647, *p* = 0.40; FA, Days: *F* (17, 86) = 2.7330, *p* = 0.0012; Groups: *F* (1, 86) = 41.5010, *p <* 10*^−^*^8^; Days*Groups: *F* (17, 86) = 0.7255, *p* = 0.77). **d**, False alarm lick rate in reinforced light-on trials (two-way ANOVA, Days: *F* (17, 86) = 0.8663, *p* = 0.6140; Groups: *F* (1, 86) = 89.3004, *p <* 10*^−^*^14^; Days*Groups: *F* (17, 86) = 3.2285, *p <* 10*^−^*^3^). **e**, False alarm lick latency in reinforced light-on trials (two-way ANOVA, Days: *F* (17, 86) = 2.0216, *p* = 0.018; Groups: *F* (1, 86) = 251.7387, *p <* 10*^−^*^26^; Days*Groups: *F* (17, 86) = 4.8600, *p <* 10*^−^*^6^). **f**, Accuracy in probe light-off trials (two-way ANOVA, Days: *F* (5, 30) = 8.3041, *p <* 10*^−^*^4^; Groups: *F* (1, 30) = 4.7288, *p* = 0.038; Days*Groups: *F* (5, 30) = 0.7288, *p* = 0.619). **g**, Action rate in probe light-off trials (two-way ANOVAs, HIT, Days: *F* (5, 30) = 5.4632, *p* = 0.0011; Groups: *F* (1, 30) = 6.3510, *p* = 0.017; Days*Groups: *F* (5, 30) = 1.2158, *p* = 0.33; FA, Days: *F* (5, 30) = 5.5019, *p* = 0.0010; Groups: *F* (1, 30) = 0, *p* = 1; Days*Groups: *F* (5, 30) = 1.1320, *p* = 0.37). **h**, HIT lick latency in probe light-off trials (two-way ANOVA, Days: *F* (5, 29) = 6.0308, *p <* 10*^−^*^3^; Groups: *F* (1, 29) = 10.3058, *p* = 0.0032; Days*Groups: *F* (5, 29) = 0.1542, *p* = 0.98). **i**, Maximal difference between hit and false alarm rates in probe light-off trials over the first 6 days (t-test, *p* = 0.40). **j**,Accuracy in reinforced light-off trials (two-way ANOVA, Days: *F* (17, 86) = 8.3579, *p <* 10*^−^*^11^; Groups: *F* (1, 86) = 1.6832, *p* = 0.20; Days*Groups: *F* (17, 86) = 0.2356, *p* = 1). **k**, Action rate in reinforced light-off trials (two-way ANOVAs, HIT, Days: *F* (17, 86) = 11.1314, *p <* 10*^−^*^14^; Groups: *F* (1, 86) = 2.1423, *p* = 0.15; Days*Groups: *F* (17, 86) = 0.9107, *p* = 0.56; FA, Days: *F* (17, 86) = 4.2760, *p <* 10*^−^*^5^; Groups: *F* (1, 86) = 0.5043, *p* = 0.48; Days*Groups: *F* (17, 86) = 0.3026, *p* = 1). **l**, FA lick latency in reinforced light-off trials (two-way ANOVA, Days: *F* (17, 78) = 1.7364, *p* = 0.053; Groups: *F* (1, 78) = 9.0848, *p* = 0.0035; Days*Groups: *F* (17, 78) = 1.3749, *p* = 0.17). **m**, FA lick rate in reinforced light-off trials (two-way ANOVA, Days: *F* (17, 78) = 0.7983, *p* = 0.69; Groups: *F* (1, 78) = 13.4564, *p <* 10*^−^*^3^; Days*Groups: *F* (17, 78) = 1.4494, *p* = 0.14). **n**, Comparison of light-off versus light-on trials to measure auditory cortex silencing effect on on-line performance. **o**, Session density plot of accuracy in reinforced light-on against light-off. Top, control; bottom, PV-ChR2. **p**, Accuracy in light-on reinforced trials from day where FA*<* 0.3 in light-off reinforced trials. Note the general trend for ChR2 mice (gray lines) to increase accuracy (positive slopes), i.e. performance impairment fades away. **q**, Within subject difference between accuracy in reinforced light-on and light-off aligned to the day where false alarm rate *<* 0.3 in reinforced light-off.

**Supplementary Figure 15.**
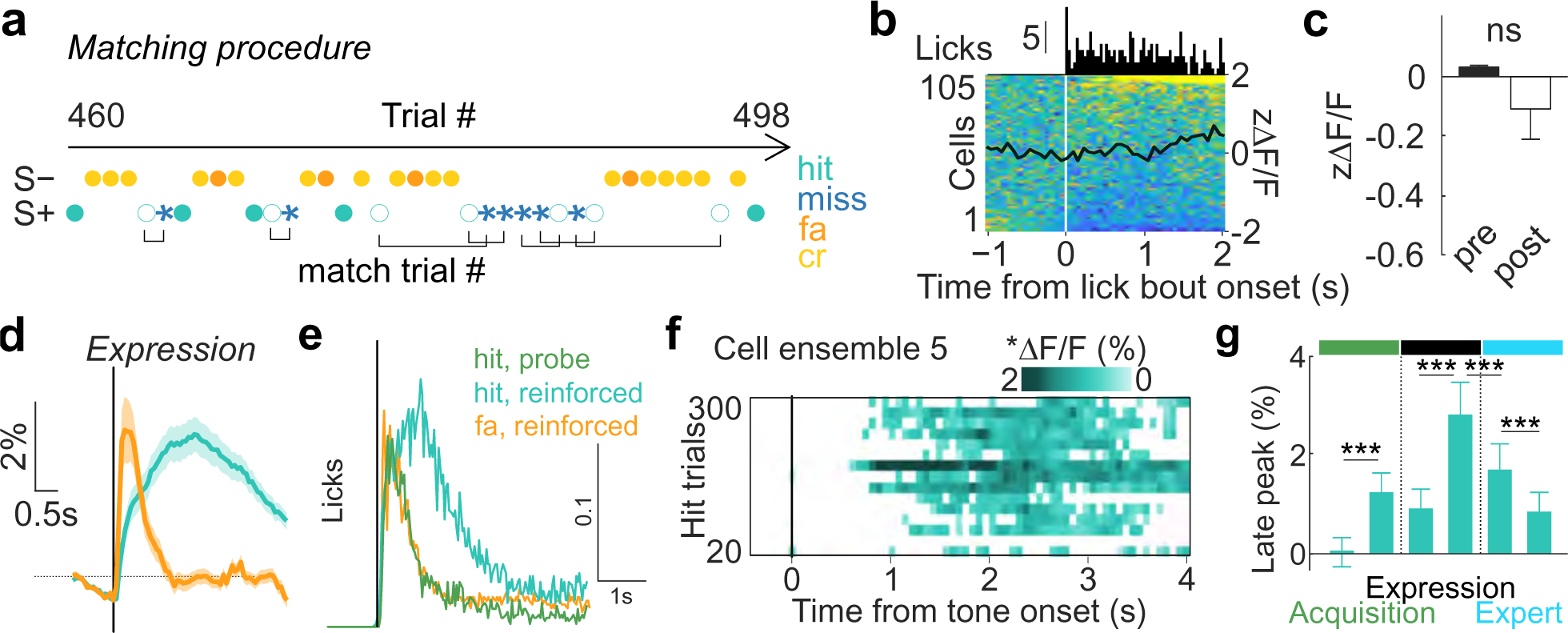
Emergence of reward prediction signal. **a**, Procedure of hit and miss trial matching. **b**, Heat map of members of cell ensemble 5 (*n* = 105) activity aligned to lick bout onset outside task events in day 1 of training. Lick PSTH is represented above. **c**, Quantification of z-scored calcium activity 1s pre- vs 1s post-lick bout onset (Wilcoxon test, *p* = 0.11). **d**, Average cell ensemble 5 activity in reinforced hit (green) and FA (orange) trials over Expression phase. **e**, Lick PSTHs aligned to tone onset of FA trials in Expression and hit trials in probe context. **f**, Cell ensemble 5 activity over the first 300 hit trials (20-trial blocks). Only significant activity (and higher than null population, see Methods) is represented. Note the emergence of a stable late-on-trial signal after 40 hit trials onwards. **g**, Quantification of Fig. 4l, i.e. evolution of late-in-trial signal of cell ensemble 5 across learning, taking first and last two 40-hit trial blocks (KW test, *p* = 1.05.10*^−^*^23^). \**p <* 0.05, \*\**p <* 0.01, \*\*\**p <* 0.001, n.s.: not significant.

**Supplementary Figure 16.**
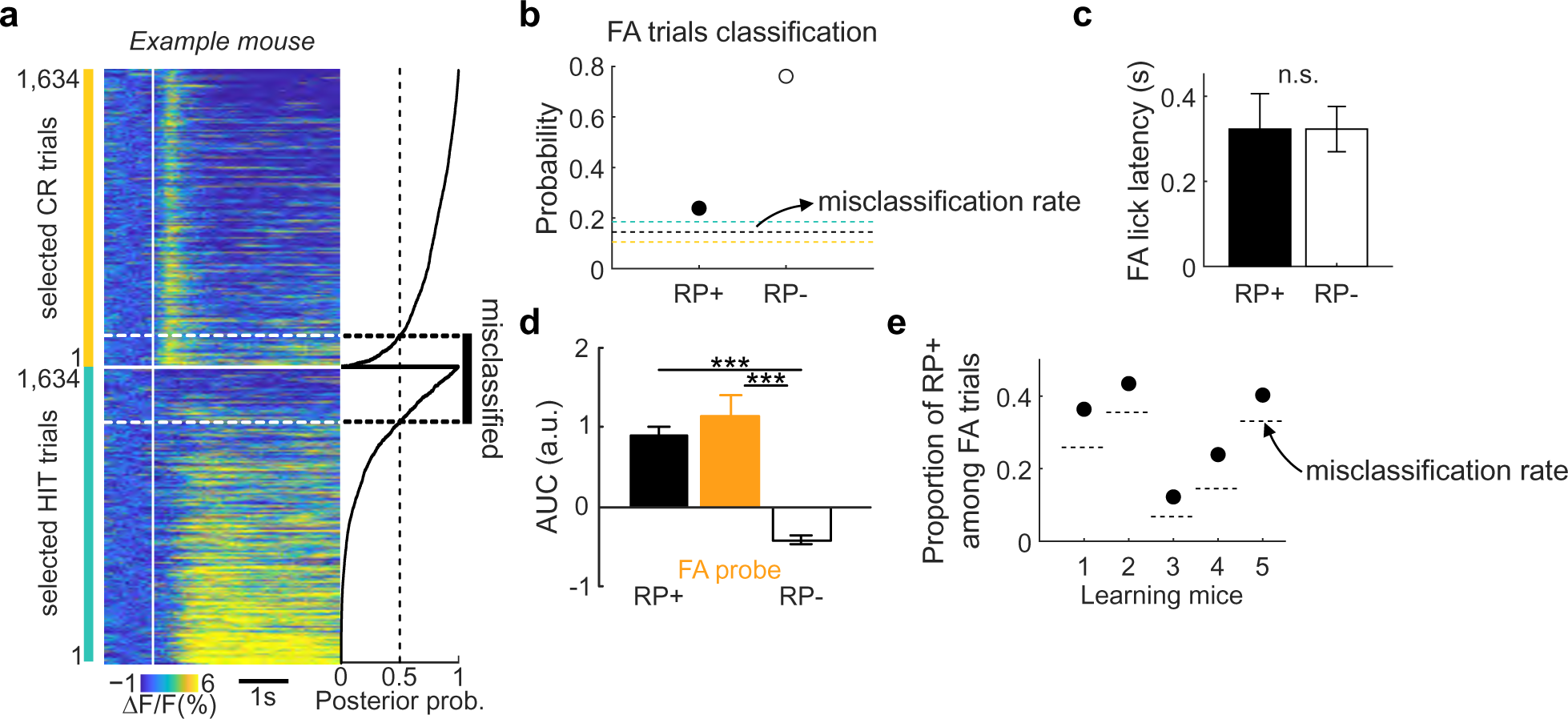
Reward prediction signal on error trials. **a**, Classification of hit versus CR trials in the reinforced context from the AUC post-tone of a fraction of cell ensemble 5 (*n* = 51) recorded in the example mouse showed in Fig. 4p,q. Right: posterior probability of being part of CR class. **b**, Proportion of RP+ and RP*−* FA trials from the example mouse showed in Fig. 4o,p. **c**, No difference in lick latency was observed between RP+ and RP*−* FA trials (Wilcoxon test, *p* = 0.83). **d**, AUC quantification of RP+, RP*−* and probe FA trials (KW, *p* = 9.76.10*^−^*^28^). **e**, Proportion of RP+ among all FA trials and misclassification rate in each learning mice. \**p <* 0.05, \*\**p <* 0.01, \*\*\**p <* 0.001, n.s.: not significant.

**Supplementary Figure 17.**
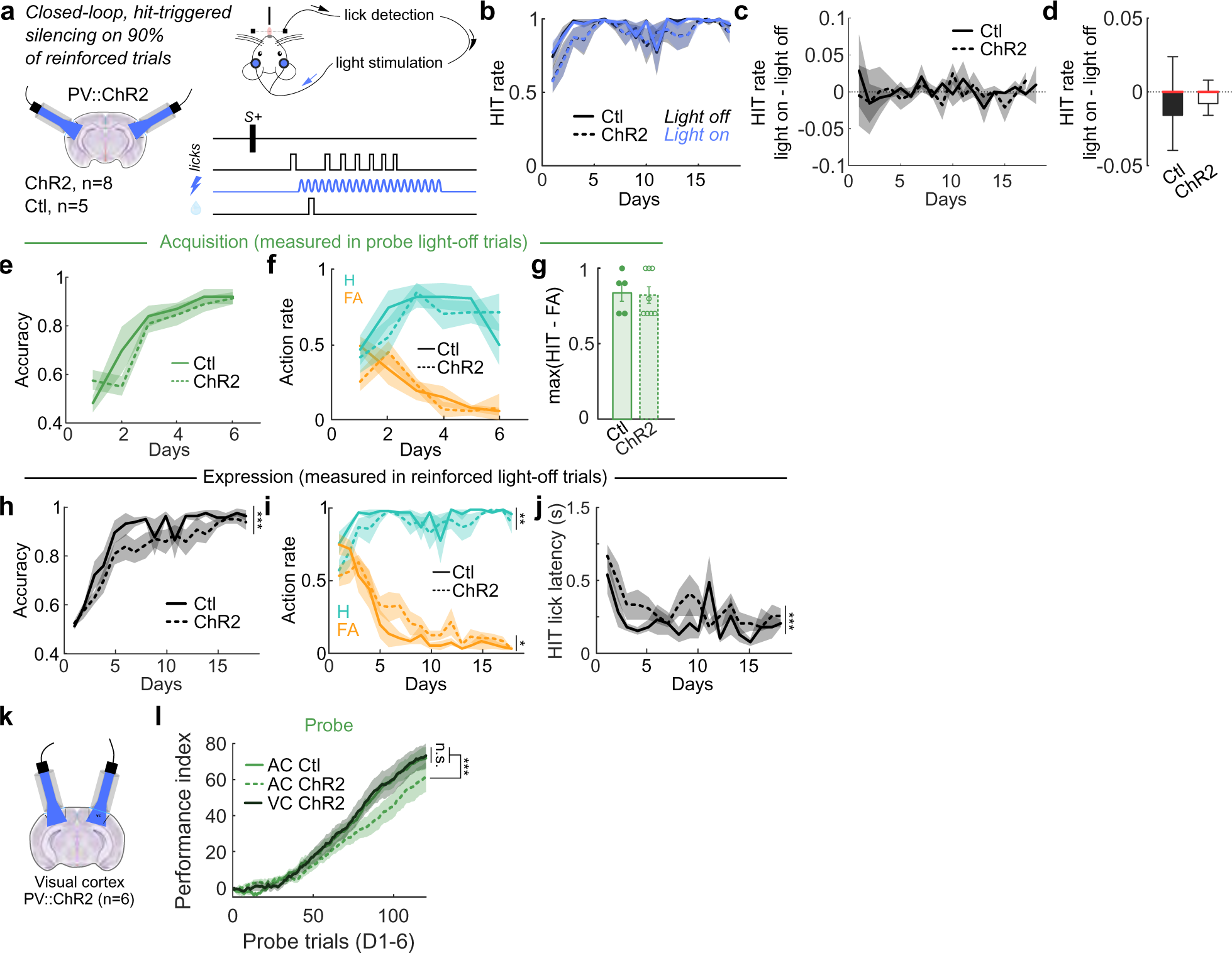
Post-hit silencing over learning. **a**, Experimental design of optogenetic silencing of AC activity throughout learning post hit only. **b**, Hit rate across days in control (Ctl) and test (ChR2) mice in reinforced light-on or light-off trials across days. **c**, Difference in hit rate in reinforced light-on versus light-off trials across days. **d**, Difference in hit rate in reinforced light-on versus light-off trials (Wilcoxon test, *p* = 0.13). **e**, Accuracy in probe light-off trials. **f**, Action rate (hit, H; false alarm, FA) in probe light-off trials. **g**, Maximum difference between hit and false alarm trials over the first 6 days in probe light-off trials. **h**, Accuracy in reinforced light-off trials. **i**, Action rate in reinforced light-off trials. **j**, Hit lick latency in reinforced light-off trials. **k**, Silencing of visual cortex (VC) activity throughout learning post hit only. **l**, Performance index in probe trials for AC control (*n* = 5), AC PV-ChR2 (*n* = 8) and VC PV-ChR2 (*n* = 6) (two-way ANOVA, *p* = 1.90.10*^−^*^32^).

**Supplementary Figure 18.**
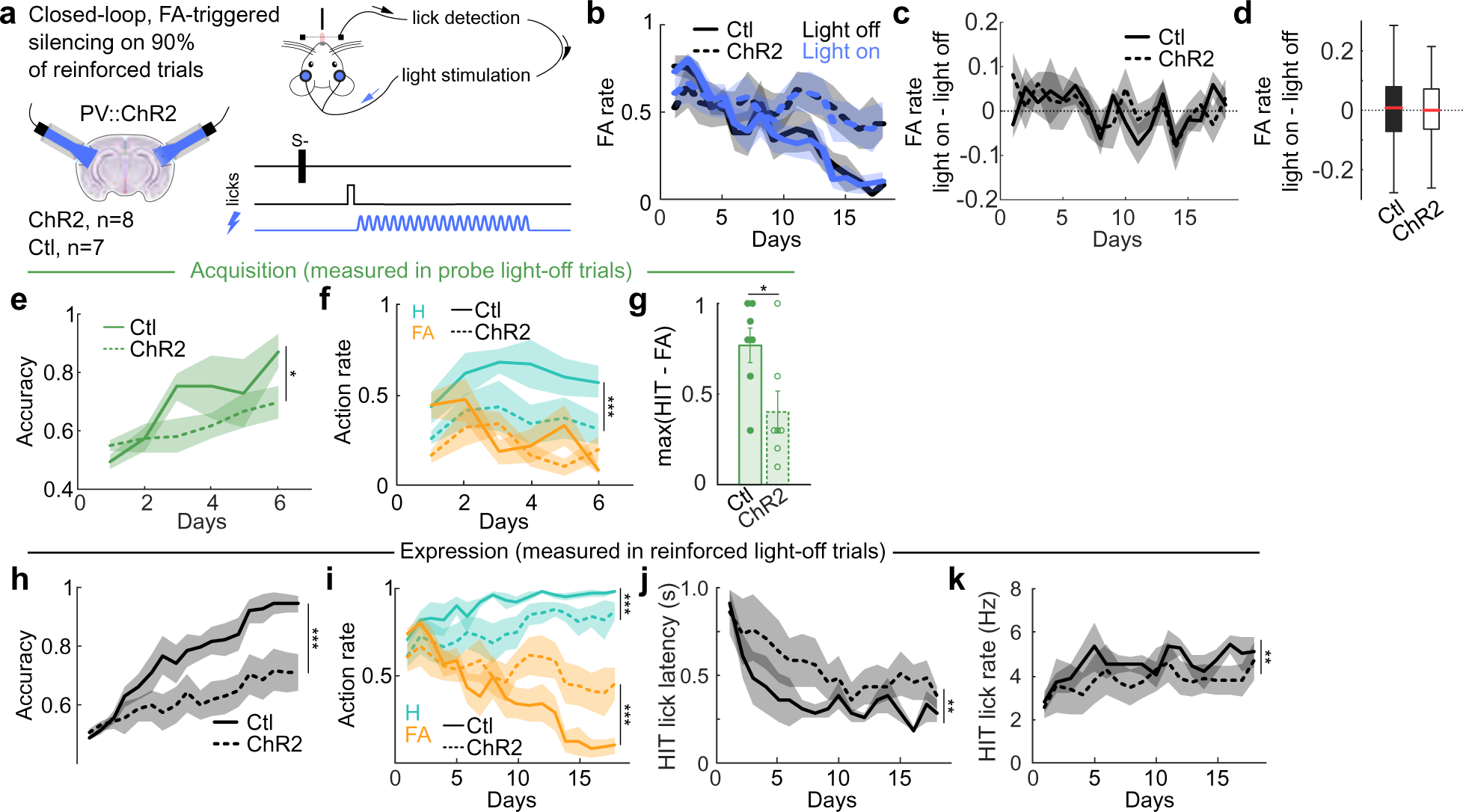
Post-FA silencing over learning. **a**, Experimental design of optogenetic silencing of AC activity throughout learning post false alarm (FA) only. **b**, False alarm rate across days in control (Ctl) and test (ChR2) mice in reinforced light-on or light-off trials across days. **c**, Difference in false alarm rate in reinforced light-on versus light-off trials across days. **d**, Difference in false alarm rate in reinforced light-on versus light-off trials (t-test, *p* = 0.76). **e**, Accuracy in probe light-off trials. **f**, Action rate (hit, H; false alarm, FA) in probe light-off trials. **g**, Maximum difference between hit and false alarm trials over the first 6 days in probe light-off trials. **h**, Accuracy in reinforced light-off trials. **i**, Action rate in reinforced light-off trials. **j**, Hit lick latency in reinforced light-off trials. **j**, Hit lick rate in reinforced light-off trials.

**Supplementary Figure 19.**
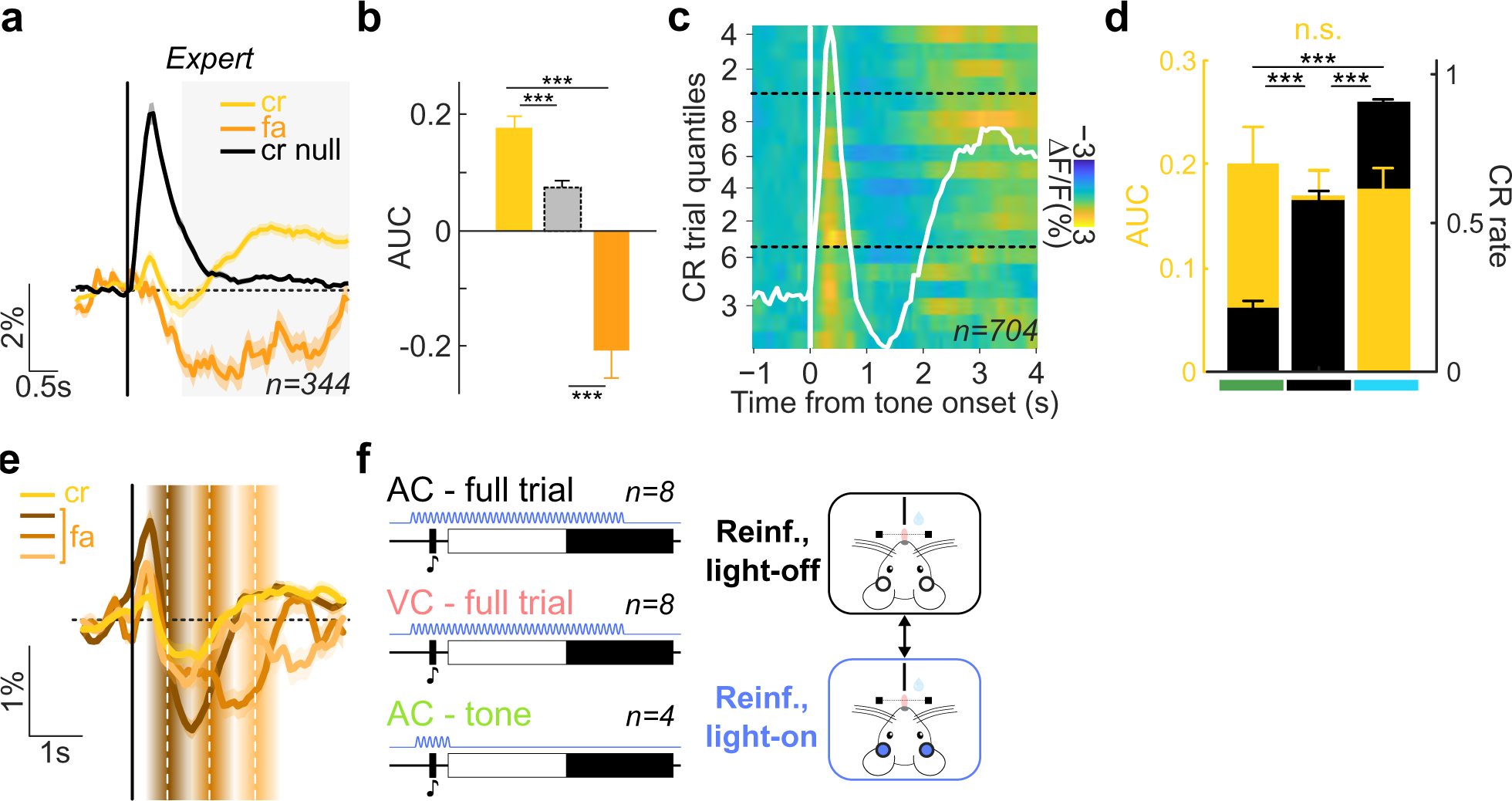
A signal for action suppression in Learning network. **a**, Average activity of cell ensemble 6 or low weighted cells (null, black) in CR and FA trials in Expert phase. **b**, Quantification of late-in-trial activity (KW test, *p* = 4.76.10*^−^*^44^). **c**, Average cell ensemble 6 activity across learning phases. CR trials were split into 6, 9 and 4 quantiles over Acquisition, Expression and Expert phases, respectively. **d**, Quantification of late-in-trial activity (left axis) and CR rate (right axis) over learning phases. **e**,Averaged ensemble 6 activity in FA and CR trials. FA trials are split according to lick latencies (white dashed line, mean latency; graded rectangles, latency range extrema). **f**, Silencing protocols compared in Fig. 5h,i.

**Supplementary Table 1.**
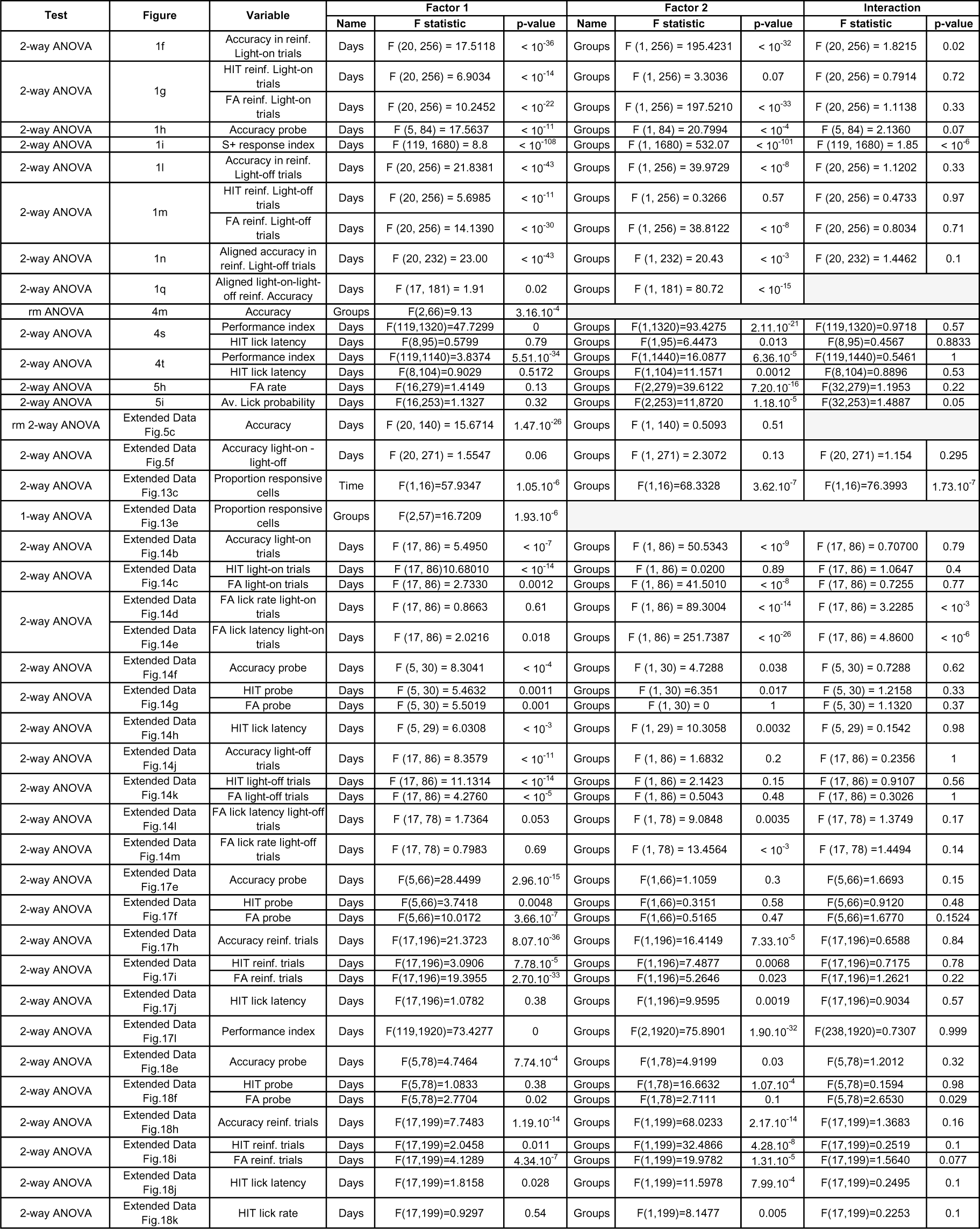
Report of ANOVA statistics.

## Notes

### Competing Interest Statement

The authors have declared no competing interest.

